# How are different contrast effects on binocular luster spatially integrated?

**DOI:** 10.64898/2026.01.06.697891

**Authors:** Gunnar Wendt, Franz Faul

## Abstract

In previous studies with dichoptic center-ring-surround stimuli, we found that two properties of the ring element have a strong influence on the phenomenon of binocular luster. The strength of the lustrous impression in the central target patch varies with increasing ring width, with the direction of this variation (increasing or decreasing) depending on the ring’s luminance. In this study, we used stimuli in which the ring was split into segments with two different luminances that in uniform rings had opposite effects on perceived luster. The aim was to investigate how the lustrous impression is influenced by combining a weaker and a stronger contrast effect, in particular how they are spatially integrated by the visual system. In a psychophysical matching experiment, subjects had to assess the strength of the lustrous impression in a series of test stimuli with different ring widths, numbers of ring segments, and spatial proportions between the two ring parts. We found that the results of the experiment could neither be explained by a winner-takes-all integration (assuming that the lustrous response is completely determined by the stronger effect) nor by a balanced integration process (assuming equal weights for the two effects). Instead, both effects contribute to the overall lustrous response, with the stronger effect having a greater weight. Interestingly, the magnitude of this weight varied considerably between different groups of subjects. We found two main trends in the data, representing two different types of sensitivity to the phenomenon of binocular luster.

## 1. Introduction

The phenomenon of binocular luster can be observed when the two eyes are presented with stimuli that differ in color or luminance contrast. For the study of this phenomenon, simple dichoptic center-surround configurations have been established as standard stimuli in which the center patches differ in color or luminance whereas the surrounds are identical (Anstis, 2000, Formankiewicz & Mollon, 2009, Jung, Moon, Park, & Song, 2013, Kiesow, 1920, Malkoc & Kingdom, 2012, Sheedy & Stocker, 1984, Wendt & Faul, 2019, Wolfe & Franzel, 1988, Zhang, 2015). With suitable contrasts, the binocularly combined center patch takes on a strange perceptual quality, usually described as shimmering or lustrous, sometimes also as fluorescent or luminous (see also Wendt & Faul, 2022b). Using this type of stimulus, Anstis (2000) demonstrated that the lustrous impression is significantly stronger when the two monocular half-images have reversed contrast polarities (*inc-dec* stimuli, where a luminance increment is binocularly paired with a luminance decrement) than when the two contrasts have equal polarities (*inc-inc* or *dec-dec* stimuli, respectively) (see also Georgeson, Wallis, Meese, & Baker, 2016, Hetley & Stine, 2019, Venkataramanan, Gawde, Hathibelagal, & Bharadwaj, 2021, Wendt & Faul, 2019).

In our own experiments on binocular luster, we used a stimulus variant in which the central patches were enclosed by identical rings (Wendt & Faul, 2020, 2022a, 2024, 2025). We found that the magnitude of the lustrous response depends on two properties of the ring element. Firstly, as can be seen in Figure 1, the luster judgments vary strongly with the width of the ring. Secondly, the shape of the resulting “luster curves” is determined by the relationship between the ring’s luminance and the luminances of the surround element and the two center patches.

**Figure 1:**
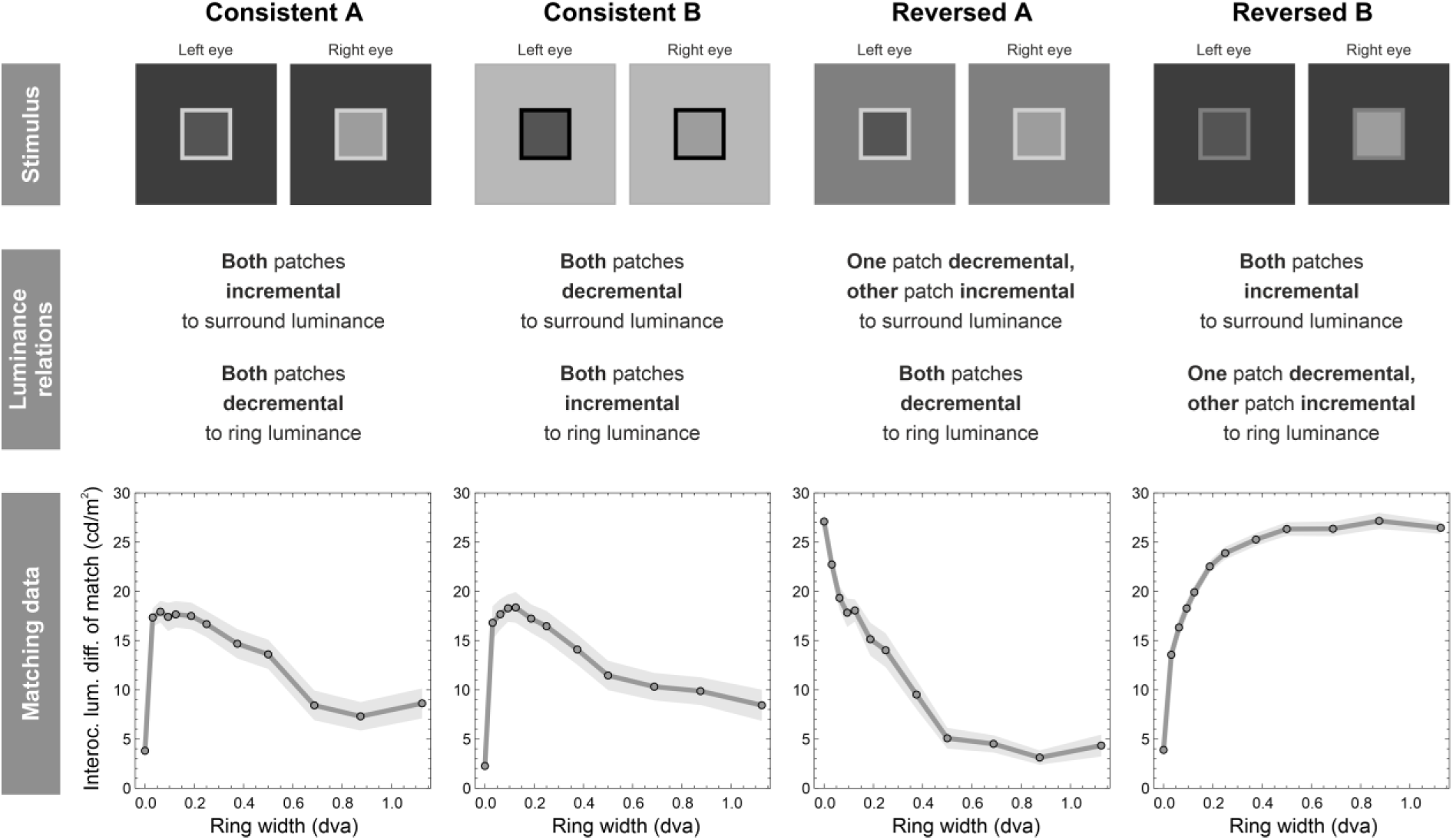
Example results from Wendt and Faul (2025). In a matching experiment, observers had to assess the strength of the lustrous impression in the central area of dichoptic center-ring-surround stimuli with varying ring widths. In addition, four different combinations of ring and surround luminance were tested. The strength of the lustrous impression varied strongly depending on the ring width. The shape of the luster curves was determined by the luminance of the ring in relation to the luminances of the surround element and the two center patches. The depicted luster settings were obtained with square center patches with a side length of 4 degrees of visual angle and a luminance difference of 30 cd/m^2^ between the monocular center patches. Transparent areas represent the SEM in both directions.

In our previous work, we have shown that all these observations can be explained by a simple neural process located at an early stage of the visual pathway (Wendt & Faul, 2022a, 2025; see also Appendix A). The basic component of this mechanism is a binocular cell that receives its input signals from two types of monocular contrast detector cells whose receptive fields have reversed excitatory and inhibitory areas, making them sensitive to light patterns with opposite polarity. Suitable candidates for such monocular cells include neurons with an antagonistic circular-symmetric center-surround organization of their receptive fields (similar to the retinal ganglion ON- and OFF-center cells; see Schiller, 1992), as well as orientation- selective cells, whose excitatory and inhibitory subregions are arranged side-by-side (for example, with a Gabor-like layout where the ON- and OFF-cells are represented by opposite phases). One main characteristic of the binocular cell is that its response strongly depends on the types of monocular cells that provide the input signals from corresponding locations of the two eyes. Signals from monocular cells of the same type (ON-ON or OFF-OFF pairings) produce much weaker responses than the pairing of an ON-mechanism with an OFF-mechanism.

The results of adaptation experiments conducted by Kingdom, Jennings, and Georgeson (2018) support the idea that the phenomenon of binocular luster is mediated by specific binocular neurons. Evidence for the existence of binocular cells with the above-mentioned response characteristics has already been provided by Poggio and Fischer (1977). These binocular neurons, located in the primary visual cortex and referred to as *tuned-inhibitory* cells, have been shown to be highly sensitive to monocular contrast signals with reversed polarity and may serve as a type of interocular conflict detector (Read & Cumming, 2007; see also Goncalves & Welchman, 2017, Katyal, Engel, He, & He, 2016, Katyal, Vergeer, He, He, & Engel, 2018). Kingdom, Read, Hibbard, and May (2022) therefore assume that these tuned-inhibitory cells may play a crucial role in the luster phenomenon.

Taking the *Consistent A* and the *Reversed B* conditions as examples (see Fig. 1), we demonstrate in Appendix A how such a mechanism can account for the existing findings (see also Wendt & Faul, 2022a, 2025). These two contrast conditions also play a central role in the present study, as they have some interesting properties: Although they differ only in the luminance of the ring element, this difference has an almost opposite effect on the luster curves. While the lustrous response in the *Reversed B* condition becomes stronger with increasing ring width, it generally decreases monotonically in the *Consistent A* condition (after an initial steep increase at very small ring widths, see Fig. 1).

In this study, we employed a ‘hybrid’ of these two contrast conditions to investigate how perceived luster depends on more complex stimulus patterns that combine different contrast effects. This can be achieved by dividing the ring into segments that differ either in luminance alone or in both luminance and width (Fig. 2). Our study specifically addresses three questions: (1) How are the different influences on the luster phenomenon spatially integrated? For example, do they contribute equally to the overall response (balanced integration) or is the response determined by the stronger of the two effects (winner-takes-all)? (2) Can the luster judgments obtained with such mixed stimuli be predicted by our interocular conflict model, which implements a balanced integration (Wendt & Faul, 2022a, 2025)? Would a circular-symmetric layout of the receptive field of the monocular contrast detector cells lead to different predictions than an orientation-selective side-by-side organization of the excitatory and inhibitory regions? (3) Finally, is the lustrous impression within the target area of these mixed stimuli spatially homogeneous, and if not, how does the perceived inhomogeneity affect the overall luster judgments?

**Figure 2:**
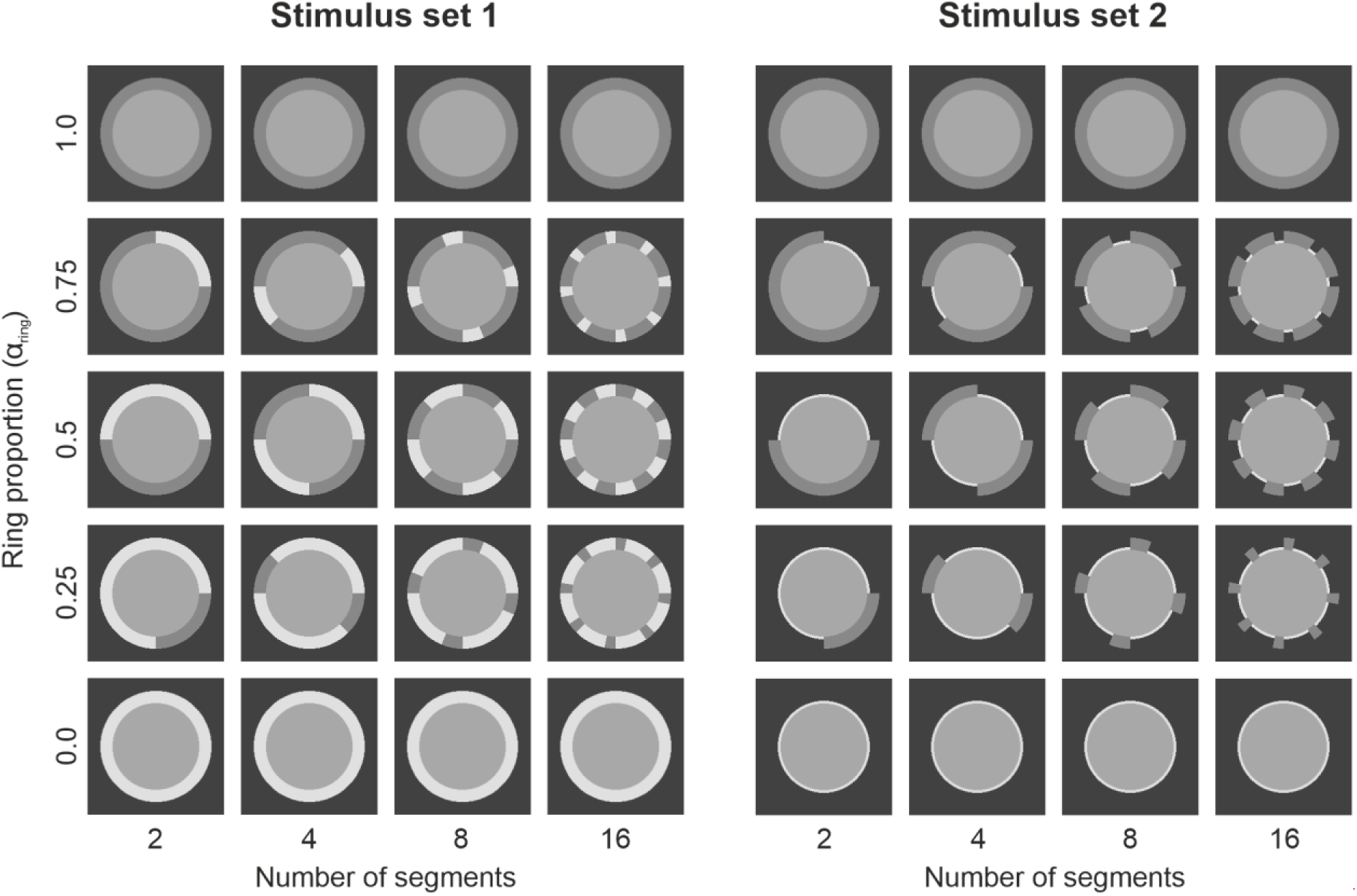
Ring structures as they were used in the two different stimulus sets. In set 1 (left panel), the two ring parts generally had different luminances (except for stimuli with ring proportions of 0.0 or 1.0, respectively) but the same ring width in each stimulus (here shown with a fixed ring width of 0.5625 dva). In set 2 (right panel), one ring part had a constant width of 0.5625 dva and a luminance of 25 cd/m^2^, whereas the other ring part had a variable width and a luminance of 75 cd/m^2^ (here shown with a constant ring width of 0.125 dva).

## 2. Methods

### 2.1. Observers

Eight subjects participated in the experiments, one of them being the first author of the study (GW). Five of the participants were female and three male, with their ages ranging from 20 to 53 years. All subjects had normal or corrected-to-normal visual acuity. Prior to the experiments, we obtained written informed consent from all subjects. The study was conducted in agreement with the ethical standards of the Deutsche Forschungsgemeinschaft (German Research Foundation, DFG) and was approved by the Central Ethics Committee of Kiel University (ZEK-18/24).

### 2.2. Stimuli and apparatus

In order to test how the visual system spatially integrates different contrast effects on binocular luster, we used dichoptic center-ring-surround configurations with a circular shape as test stimuli that were mixed versions of the two original contrast conditions *Consistent A* and *Reversed B* (see Fig. 1). We used stimuli in which the ring element was subdivided into either 2, 4, 8, or 16 segments, where neighboring segments either differed in luminance only (stimulus set 1, left panel in Fig. 2), or both in luminance and width (stimulus set 2, right panel in Fig. 2). In addition, we varied the relative length of the two segment types. The ring proportions are defined by the mixing factor *α_ring_*, which represents the relative length of the stronger ring part (based on the *Reversed B* contrast conditions, see Fig. 1), whereas the weaker part (based on the *Consistent A* condition) has a proportion of 1- *α_ring_*. Stimuli with *α_ring_* = 0, 0.25, 0.5, 0.75, and 1.0 were tested. Stimuli with *α*_ring_ = 0 and 1.0 are the two original contrast conditions with a uniform ring. Finally, we varied the width of the ring segments, either equally for the two ring parts (stimulus set 1), or for one ring part only, while the other ring part was kept at a constant width (stimulus set 2). Ring widths of 0.0, 0.0625, 0.125, 0.25, 0.375, 0.5625, 0.8125, and 1.0625 dva were used for the varying part of the ring in stimulus set 2. In stimulus set 1, however, the no-ring condition (0 dva) was omitted, because this condition would produce identical center-surround stimuli for all combinations of segment number and ring proportion. The luminance of the common surround, which covered the entire upper half of the monitor in which the test stimuli were presented, always had a constant luminance of 5 cd/m^2^. The circular center patches, representing the target area of the stimuli, had a radius of 2.0 dva and a luminance of 10 cd/m^2^ for one eye and 40 cd/m^2^ for the other (these luminances were swapped between eyes in half of the trials to account for potential imbalances in eye dominance).

The stimuli were displayed on a monitor with a diagonal of 24 inches (EIZO CG243W) and a resolution of 1920 × 1200 pixels. To color calibrate the monitor, a JETI specbos 1211 spectroradiometer was used, following a method described in Brainard (1989). The dichoptic stimulus pairs were presented side by side on the screen and fused using a mirror-stereoscope (ScreenScope) mounted on the monitor. The light paths from the screen to the observer’s eye had a length of 50 cm.

### 2.3. Procedure

In each trial, observers evaluated three properties of the test stimulus. First, they had to determine whether the target area of the test stimulus appeared lustrous at all. To this end, the test stimulus (in the top half of the monitor) was presented together with an anchor stimulus (in the bottom half). While the ring structure and the background of the anchor stimulus were always identical to those of the test, the center patches of the anchor always had the same luminance of 22.5 cd/m^2^ and thus appeared matte. This luminance was chosen such that (a) it was close to the mean luminance of the two center patches of the test stimulus (which was always 25 cd/m^2^) and (b) it still allowed for spatial discrimination between the center patch area and the ring area in cases where the ring element had a luminance of 25 cd/m^2^. This anchor stimulus was meant to facilitate the detection of weaker lustrous sensations in the test. Using the arrow keys on the keyboard, the observer could switch between the two response alternatives “lustrous” and “non-lustrous”. If „non-lustrous“ was selected, the trial was terminated and the next trial began after a dark adaptation period of three seconds. If „lustrous“ was selected, the trial proceeded to the second part, in which the test stimulus was presented together with a matching stimulus consisting of a dichoptic center-ring-surround configuration with a square outline (Fig. 3). The center patches of the match had a side-length of 2.0 dva and were enclosed by a 0.125 dva wide ring with a fixed luminance of 25 cd/m^2^.

**Figure 3:**
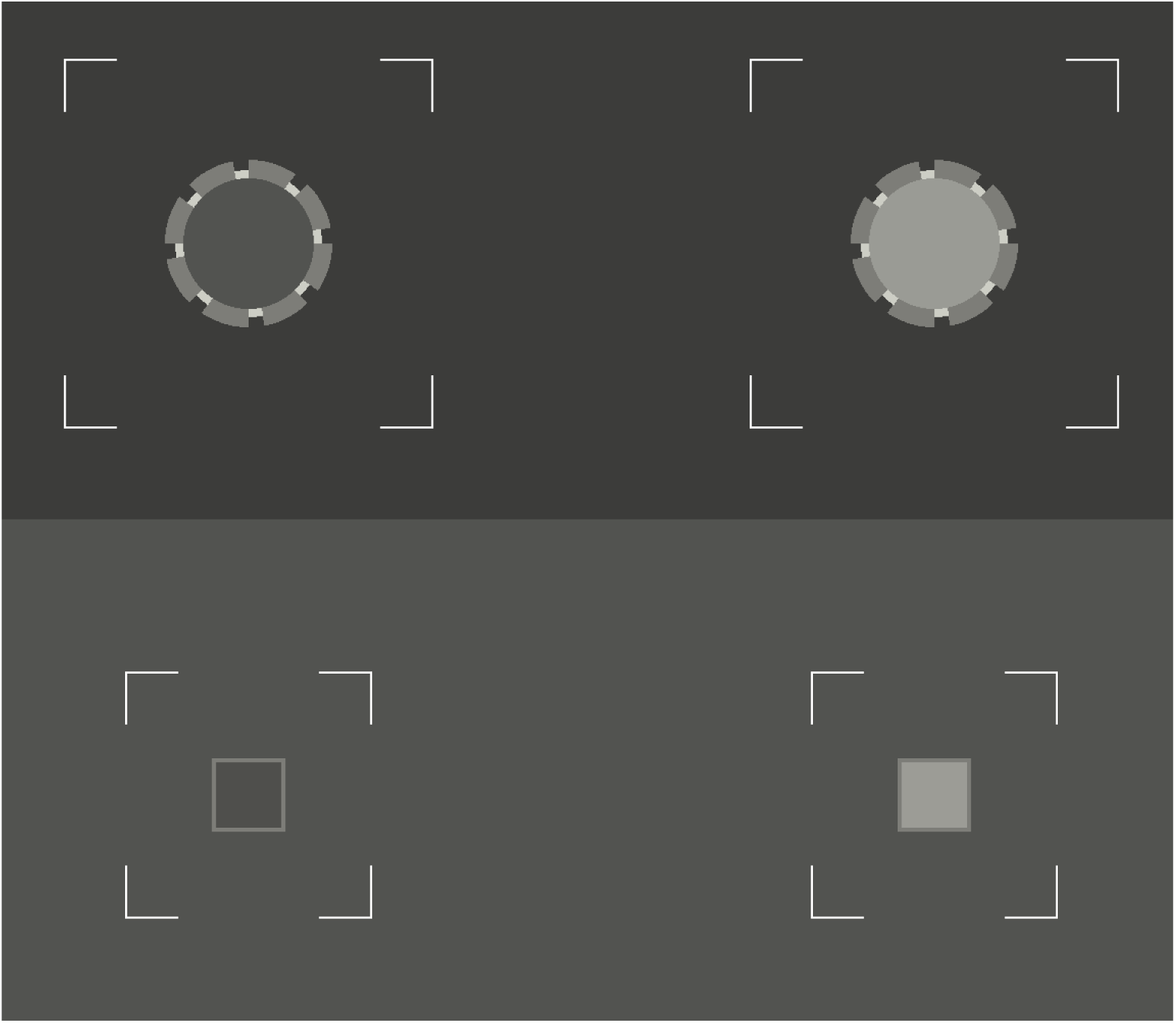
Central part of the screen area during the matching task. The dichoptic test stimulus was always presented in the upper half of the screen, the matching stimulus always in the lower half. The task of the subject was to adjust the interocular luminance difference between the two center patches of the match such that the perceived luster was indistinguishable from that of the test stimulus.

The luminance of the common surround, which filled the entire bottom half of the screen, was 10 cd/m^2^. Using the arrow keys on the keyboard, the observer had to adjust the interocular luminance difference |*C_l_ – C_r_*| between the two center patches such that the perceived luster in the match was indistinguishable from that of the target patch of the test stimulus. This luminance difference was used as a measure for the perceived luster strength (see also Wendt & Faul, 2025). This choice is supported by the strong correlations between the interocular luminance difference |*C_l_ – C_r_*| and corresponding rating values in our previous studies (Wendt & Faul, 2020, 2022a, 2024), in which the observers additionally rated the strength of the lustrous impression on a scale from 0 to 5. The observers adjusted a parameter *p* from which the two center patch luminances *C_l_* and *C_r_* of the match were determined as follows:

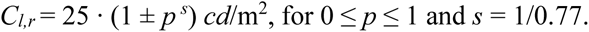

The constant *s* in this equation was selected such that the relationship between *p* and the strength of perceived luster was approximately linear (see Wendt & Faul, 2019).

Since the two different ring parts of the mixed-ring stimuli have different effects on the lustrous impression, it is possible that the perceived strength of the luster varies spatially within the target patch. For instance, areas close to the ring parts with the stronger effect could be perceived as more lustrous than areas adjacent to the weaker ring parts. This could be problematic for the matching task, as the observer has to match the perceived luster in the test with a match stimulus that always produces a uniform lustrous appearance. In these cases, a perfect match would be difficult to achieve and the observer may then use an alternative (conscious or unconscious) strategy to complete this task, for instance, by basing the luster judgment exclusively on those areas with the stronger luster or by using a weighted mixture. To check whether such effects could play a role in our study, we asked the observers in the third part of a trial to assess the perceived spatial homogeneity of the luster within the target area. To this end, the same test stimulus as in the previous two parts of the trial was shown in isolation and the observer had to select an integer rating value between 0 (= “extremely inhomogeneous”) and 10 (= “perfectly homogeneous”) using the arrow keys on the keyboard.

Prior to the experiment, each observer was carefully instructed and a series of test trials had to be performed under the supervision of the experimenter. The stimuli used in these test trials were selected from the entire set of stimulus conditions used in the experiment and provided a wide range of different luster impressions. In total, 210 different stimulus combinations were tested, each of them with four repetitions. The entire set of trials was presented in random order.

### 2.4. Calculating the predictions for the mixed-ring conditions

The empirical luster curves were evaluated with respect to two alternative integration mechanisms: a winner-takes-all integration and a balanced integration.

#### Winner-takes-all

Under a winner-takes-all integration model, the magnitudes of the lustrous responses obtained with the mixed-ring stimuli (combining a stronger and a weaker effect) should equal those of the “pure” condition with the stronger effects. That is, in combinations of the stronger *Reversed B* condition and the weaker *Consistent A* condition, a winner-takes-all mechanism would produce the same lustrous response as *Reversed B* stimuli with a homogeneous ring (with α_ring_ = 1.0, see Fig. 2). Therefore, the luster curves associated with the mixed-ring stimuli should then be identical to the curve observed with the stronger effect in the pure condition.

#### Balanced integration

In a balanced integration model, different predictions for the lustrous responses obtained with the mixed-ring stimuli can be derived. We tested two alternatives:

_(1)_ A very intuitive method is to predict these luster curves on the basis of the two pure luster curves with uniform rings. It would be reasonable to assume that the luster curve for the mixed-ring stimuli results from a weighted combination of these two curves, with the weights corresponding to the respective ring proportions. That is, with a fixed proportion of α_ring_ it would be a convex combination of the luster curve for the stronger effect weighted by α_ring_ and the luster curve for the weaker effect weighted by 1- α_ring._
(2) As an alternative to this descriptive or combined-data method for predicting the luster curves, we applied a method based on our interocular conflict model of binocular luster (Wendt & Faul, 2022a; a detailed description of the model is provided in Appendix A). Since this model incorporates a balanced integration of the conflict signals within the target area, it seems well suited for predicting the luster curves for the mixed-ring conditions. The model was fed with the same stimuli used in the experiment to predict the corresponding empirical luster curves for the mixed-ring stimuli. The model parameters required for calculating these predictions were estimated using the empirical data obtained under the two pure contrast conditions *Consistent A* and *Reversed B*. Furthermore, as mentioned in the Introduction, we used two different variants of the model for the predictions, which differ in the type of filter kernel that represents the monocular contrast detector cells. The receptive field has either a circular-symmetric center-surround organization, which was simulated with a Laplacian of Gaussian filter kernel (LoG, see Appendix A), or a side-by-side arrangement of the excitatory and inhibitory areas simulated with Gabor filters (see Appendix B). There is evidence from neurophysiological studies for the existence of both cell types in the primary visual cortex (Hubel & Wiesel, 1962; Sincich & Blasdel, 2001), making both suitable candidates for representing the monocular level in our model. The comparison between these two types of filter kernels was also motivated by the fact that mixed-ring stimuli often show a complex pattern of luminance edges that occur not only between the target area and the ring element but also between neighboring ring segments (which produce so-called T-junctions with luminance edges that are orthogonal to each other, see Fig. 2). This suggests that an orientation-selective filter kernel may lead to different predictions than a filter kernel based on an LoG kernel—and, if true, this may provide some deeper insights into the specific types of monocular contrast detector cells involved in the underlying mechanism. Since both models are based on the between-eye interaction of ON and OFF contrast mechanisms, we will refer to the two variants in the following as the LoG ON-OFF model and the oriented ON-OFF model, respectively.

## 2.5. Results

Since all mixed-ring stimuli combine the pure contrast conditions *Consistent A* and *Reversed B*, we will first take a look at the luster curves for these basic stimulus sets. The luster settings averaged across all eight subjects are shown in the top row of Figure 4 as colored symbols (left diagram, yellow for the *Consistent A* and blue for the *Reversed B* condition). Also shown are the corresponding predictions and best-fitting parameter values of the LoG (middle diagram, see also Appendix A) and the oriented ON-OFF model (right diagram, see also Appendix B). The luster curve for the *Reversed B* condition (blue) is very similar to those observed in our previous studies (see Fig. 1). The curve *for the Consistent A* condition, however, deviates considerably from previous results. The general shape of the curve is similar: it is non-monotonic with a steep increase between the no-ring condition and a ring width of 0.0625 dva, and a smoother decrease at larger ring widths. However, the peak of the curve is shifted to a much larger ring width (at 0.375 dva) and the decrease of the luster settings for subsequent ring widths is much smoother than we previously found.

**Figure 4:**
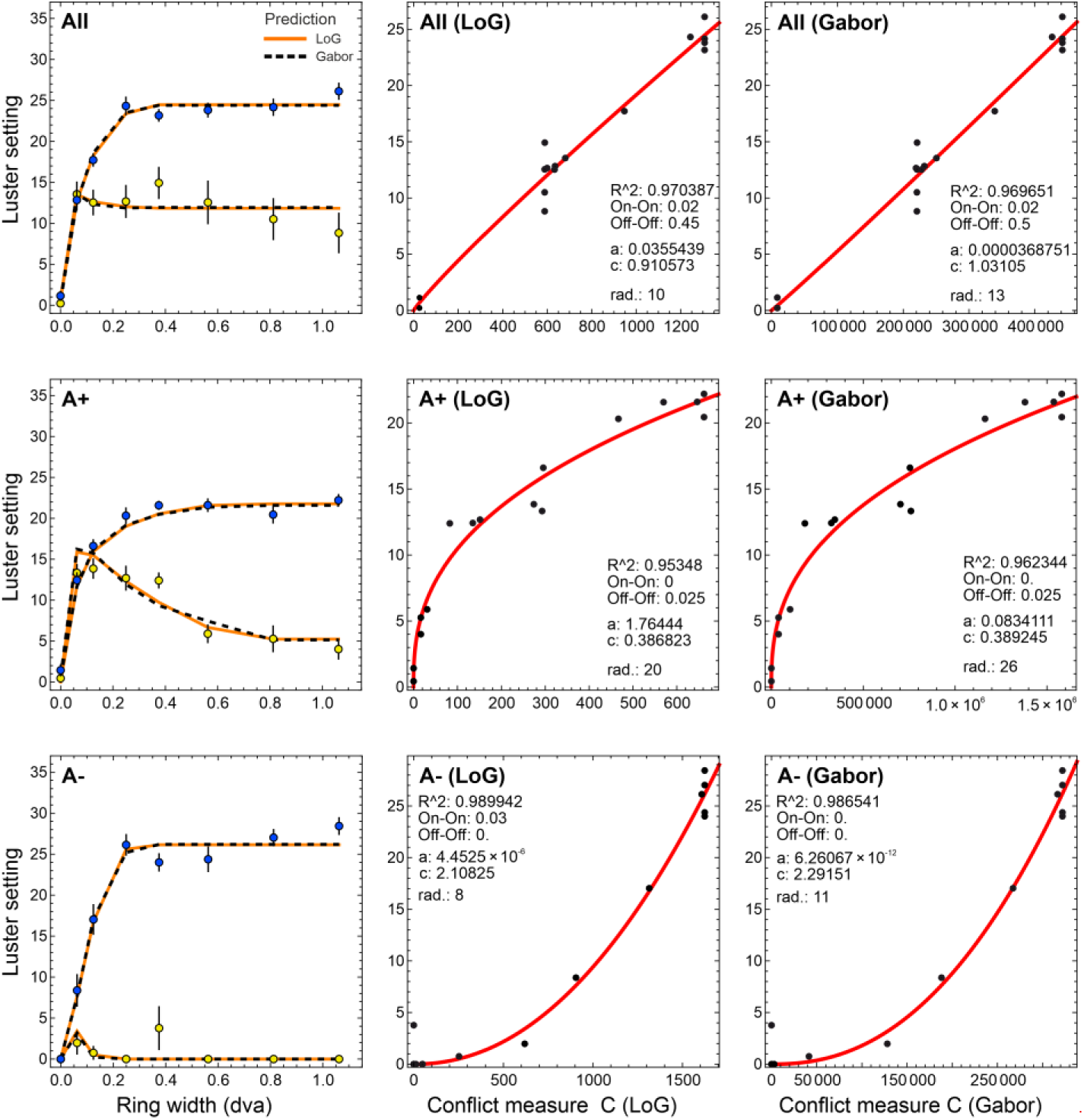
Left column: Empirical luster settings for the pure *Consistent A* (yellow disks, error bars represent the SEM in both directions) and *Reversed B* (blue disks) contrast conditions and their corresponding predictions based on the LoG (orange lines) and the oriented ON-OFF model (dashed black lines). The middle and the right diagrams show the relationship between the empirical luster settings and the conflict measure C for the LoG (middle diagram) and the oriented ON-OFF model (right diagram), respectively, based on the parameters of the best fit. Note that conflict measure C is not the final output of the model (which is C’, as it was used for the prediction lines in the left diagrams, s. also Appendix A), but an intermediate result which we use here to demonstrate the shape of the transducer functions (red curves). The top row shows the results for the entire sample of eight subjects, the middle and bottom rows those for the two sub-groups **A+** and **A-**, respectively.

The reason for this unexpected shape of the *Consistent A* luster curve becomes clearer when comparing the empirical luster settings of the individual subjects (see Appendix C). In the *Reversed B* condition, the luster curves of all subjects have a similar shape: The lustrous response increases sharply with increasing ring width until it reaches an asymptote, with slight variations in the steepness of the increase and the absolute level of the asymptote.

However, the subjects show clear differences in their sensitivity to luster in the *Consistent A* condition. There appear to be two main trends, which are also reflected in the parameter values of the corresponding model fits. One group of subjects (S01, S02, and S04), who showed high luster sensitivity for stimuli from the *Consistent A* condition (which we will refer to as the **A+** group, see the middle row in Fig. 4), produced luster curves very similar to those we have found in our previous studies. For this group, the best fit was achieved with a large radius of the filter kernel (20 and 26 pixels for the LoG and the Gabor kernel, respectively) and a nonlinear decelerating transducer function (see the red curves in the respective diagrams in Fig. 4). The other group (referred to as **A-**, see the bottom row in Fig. 4), which includes subjects S03, S06, and S07, turns out to be nearly insensitive to stimuli from the *Consistent A* condition. For this group, the best fit was achieved with a comparatively small radius of the filter kernel (8 and 11 pixels for the LoG and the Gabor kernel, respectively) and an accelerating transducer function. The two remaining subjects S05 and S08, although showing a high sensitivity for both stimulus sets, cannot be clearly assigned to either of these two groups as they produced rather unusual and, at least partly, somewhat unsystematic luster curves for the *Consistent A* condition, in combination with comparatively large standard errors. Interestingly, despite large differences in the shapes of the luster curves and corresponding differences in the model parameters, the predictive quality of the model remains very high for both the LoG and the Gabor variant (with R^2^ values for the individual datasets between 0.927 and 0.99, except for subject S08 with an R^2^ of about 0.85). Since groups **A+** and **A-** appear to indicate different types of sensitivity to the luster phenomenon, we will conduct our further analyses separately for these two groups.

Figures 5 and 6 show the empirical results of the matching task for the **A+** and the **A-** group, respectively. For each of the two stimulus sets (see the top and bottom row in each figure), the results are shown separately for the four different numbers of ring segments (columns). Each diagram shows the luster curves for the three mixed-ring conditions with the different colors representing the different proportions of the two ring parts. In addition, the two curves corresponding to the pure ring conditions are also added to each diagram in a row (see the blue and yellow curves).

**Figure 5:**
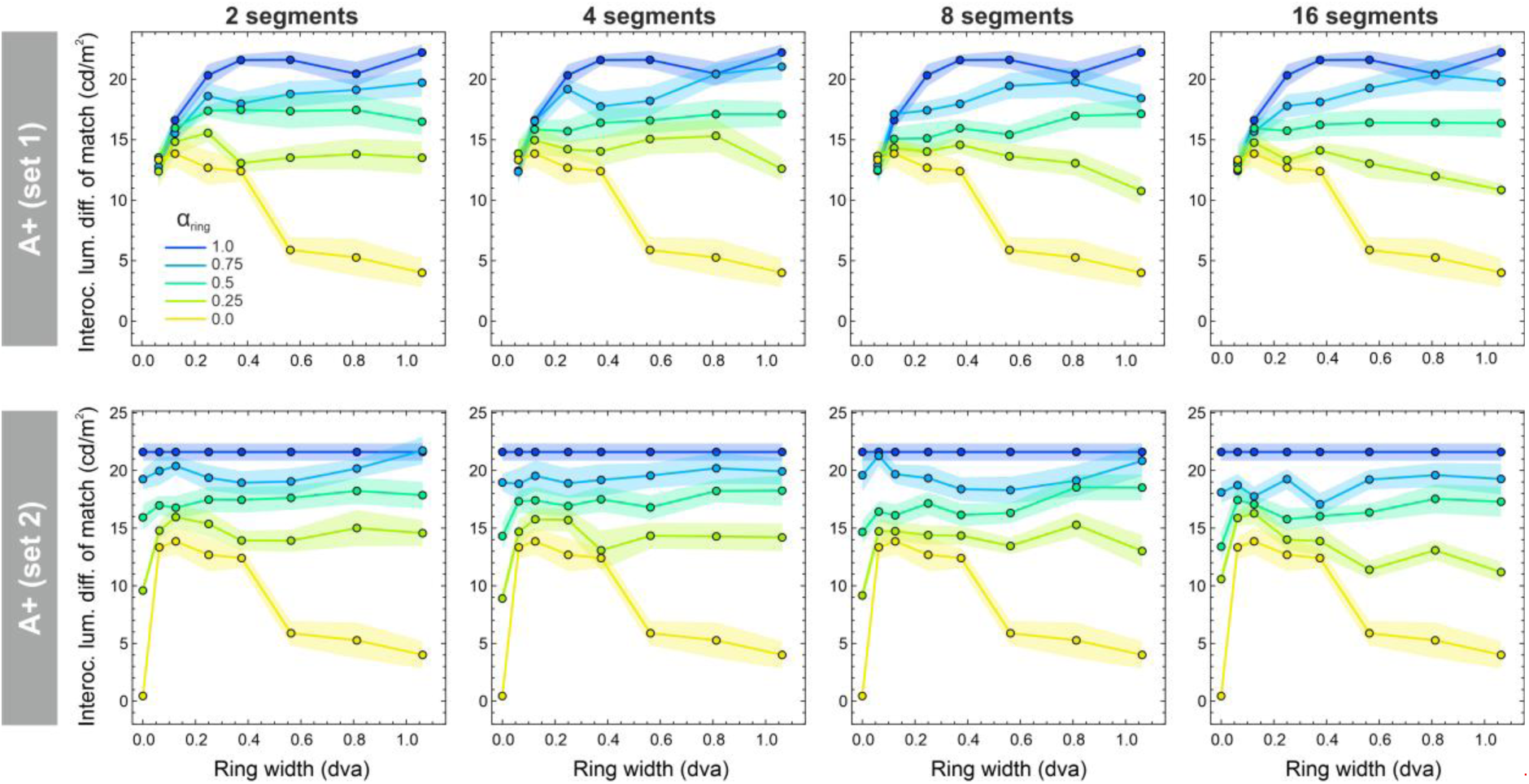
Results of the matching task of group **A+**. For each of the two stimulus sets (top and bottom row), the empirical luster settings are shown as a function of ring width, separately for the different segment numbers (columns) and the different spatial proportions for two ring parts (see the differently colored lines in each diagram; the transparent areas represent the SEM in both directions).

**Figure 6:**
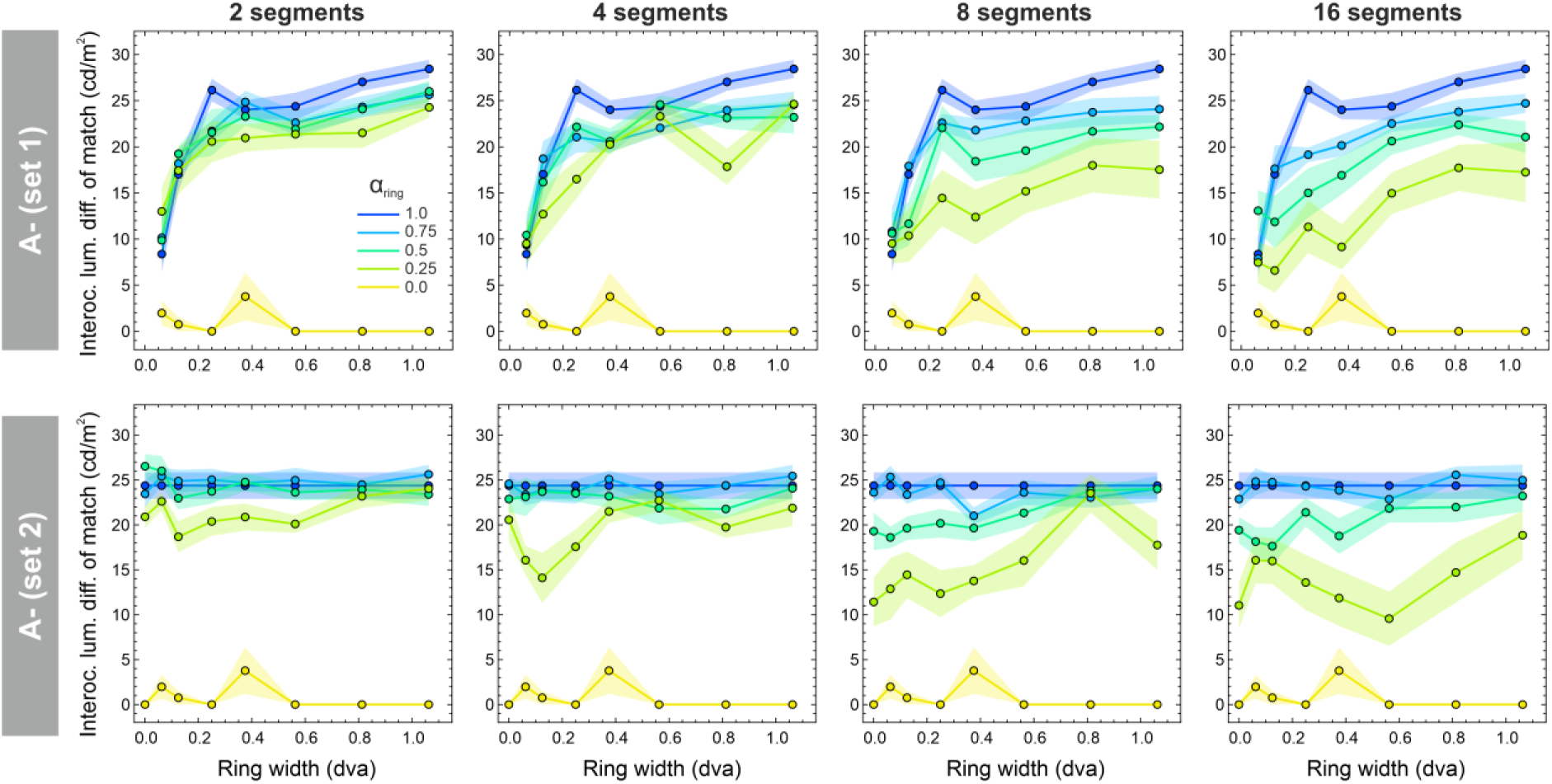
Results of the matching task of group **A-**. For each of the two stimulus sets (top and bottom row), the empirical luster settings are shown as a function of ring width, separately for the different segment numbers (columns) and the different spatial proportions for two ring parts (see the differently colored lines in each diagram; the transparent areas represent the SEM in both directions).

We will now compare the findings of both groups, starting with the results of the **A+** group. In Figure 5, one can see that the luster curves of the mixed-ring conditions lie between the two curves representing the pure contrast conditions. The order of these curves is in the expected direction, that is, the higher the spatial proportion of the ring part with the stronger effect, the more lustrous the stimulus appears. One can also immediately see that the data cannot be explained by a strict winner-takes-all mechanism. Otherwise, all luster curves for these conditions would have to be at the same level as the blue curve, which always represents the ring part with the stronger contrast effect. The luster curves of the **A-** group (Fig. 6) have the same, expected order. However, compared to those of the **A+** group, these curves much stronger tend to the blue curve, particularly at lower segment numbers.

Generally, the luster curves of the mixed-ring conditions show a strong dependency on the number of segments, an effect that is only faintly discernible in the data of the **A+** group: As the number of ring segments increases, the luster curves shift increasingly downward, i.e., the more fragmented the ring is, the weaker the lustrous impression becomes. In the discussion section, we will give two alternative explanations for this effect.

So far, it is obvious that the data cannot be explained by the assumption that the stronger of the two contrast effects fully determines the lustrous appearance in the mixed-ring stimuli (winner-takes-all). But how well are the empirical luster curves predicted by the alternative integration process, which assumes equal weights for the two effects? In Figure 7, we give a first impression of how the data of the two different subject groups **A+** and **A-** match with the corresponding predictions, as they were calculated on the basis of the LoG ON-OFF model. In Appendix D, we show the full set of prediction curves for the three different methods (that is, the combined-data as well as the two different model-based methods) together with corresponding empirical luster settings, separately for group **A+** (Fig. D1) and **A-** (Fig. D2). In the present example in Figure 7, one can see that, in general, the empirical settings for the mixed-ring conditions are above their corresponding predictions—a trend that is even more pronounced for group **A-** than for group **A+**. This means that also the alternative integration method, that is, the balanced integration, fails to explain the lustrous appearances in the mixed-ring stimuli.

**Figure 7:**
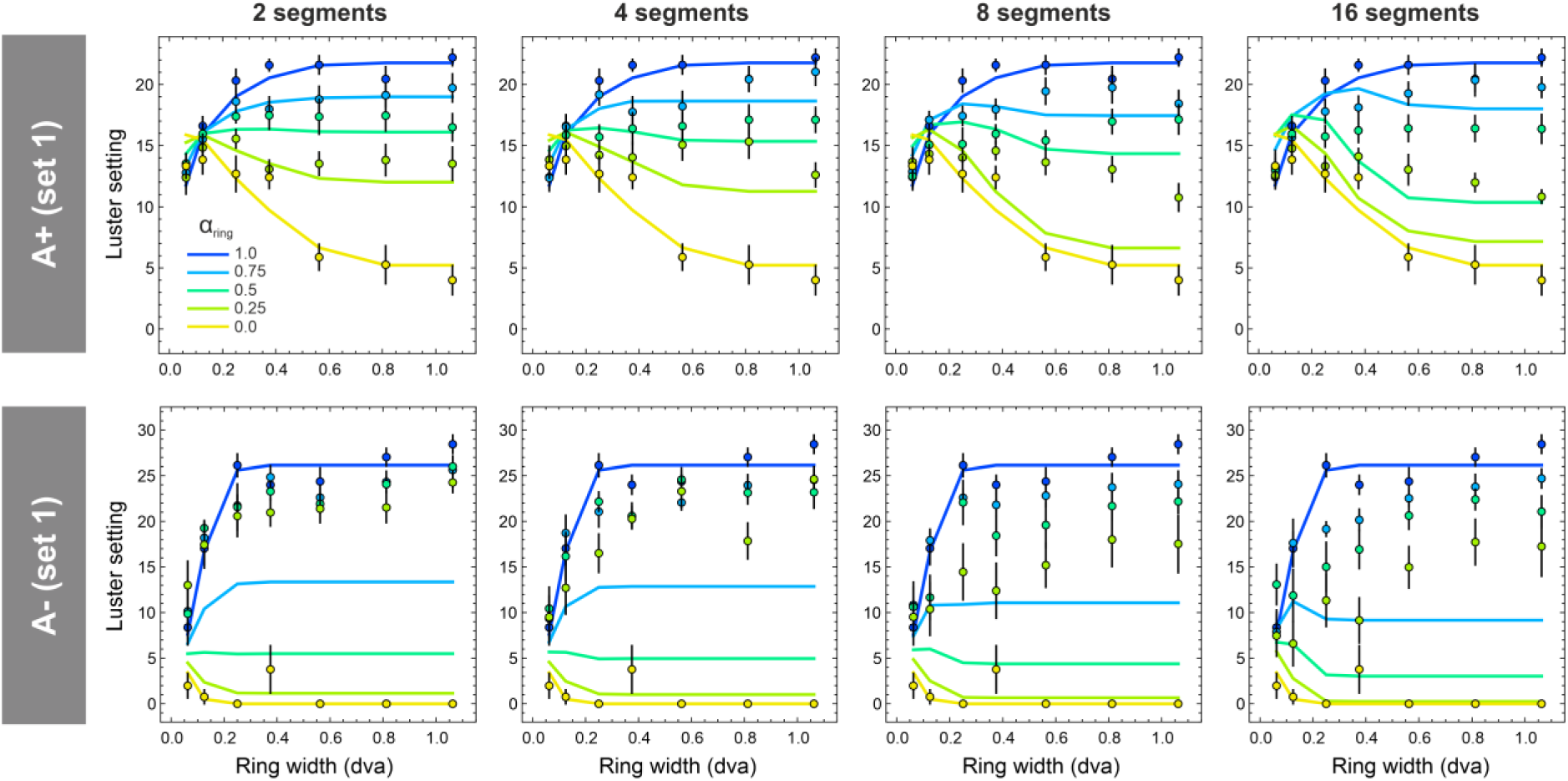
Example luster settings (colored disks) obtained with stimulus set 1 for groups **A+** (top row) and **A-**(bottom row) and their corresponding predictions (colored lines) based on the LoG ON-OFF model, separately for the different segment numbers (columns) and the different ring proportions. Generally, the empirical data points for the mixed-ring conditions (with ring proportions of 0.25, 0.5, and 0.75) lie clearly above their predictions.

To provide a better overview of how the empirical luster curves relate to the corresponding predictions for the two alternative integration methods, we calculated an index *k* that indicates how strongly the luster settings deviate from the predictions for balanced integration, relative to the predictions for the winner-takes-all integration. This means that index *k* lies in an interval between 0.0 and 1.0, where a value of 0.0 means that the given luster curve is fully consistent with the assumption of balanced integration, whereas a value of 1.0 means that the data perfectly aligns with a winner-takes-all integration process. Values between these poles mean that the two different contrast effects are integrated by an “unbalanced” mechanism in which the stronger contrast effect has a greater weight than the weaker effect, i.e., the higher the index *k*, the greater the weight for the stronger contrast effect.

For the combined-data method, in which each prediction curve is a convex mixture of the two pure luster curves with a spatially uniform ring (the blue and the yellow curves in the respective diagrams, see section 2.4), we determined the mixing factor *α_emp_* that best fits the empirical luster curve. We then set *α_emp_* in relation to the two values representing the two different integration methods, where *α_bal_* represents balanced integration (which is identical to the spatial proportion *α_ring_* of the mixed-ring stimuli) and *α_win_ =* 1.0 represents winner-takes- all integration (see the left diagram in Fig. 8): *k_data_ = (α_emp_ – a_bal_) / (1.0 – α_bal_).*

**Figure 8:**
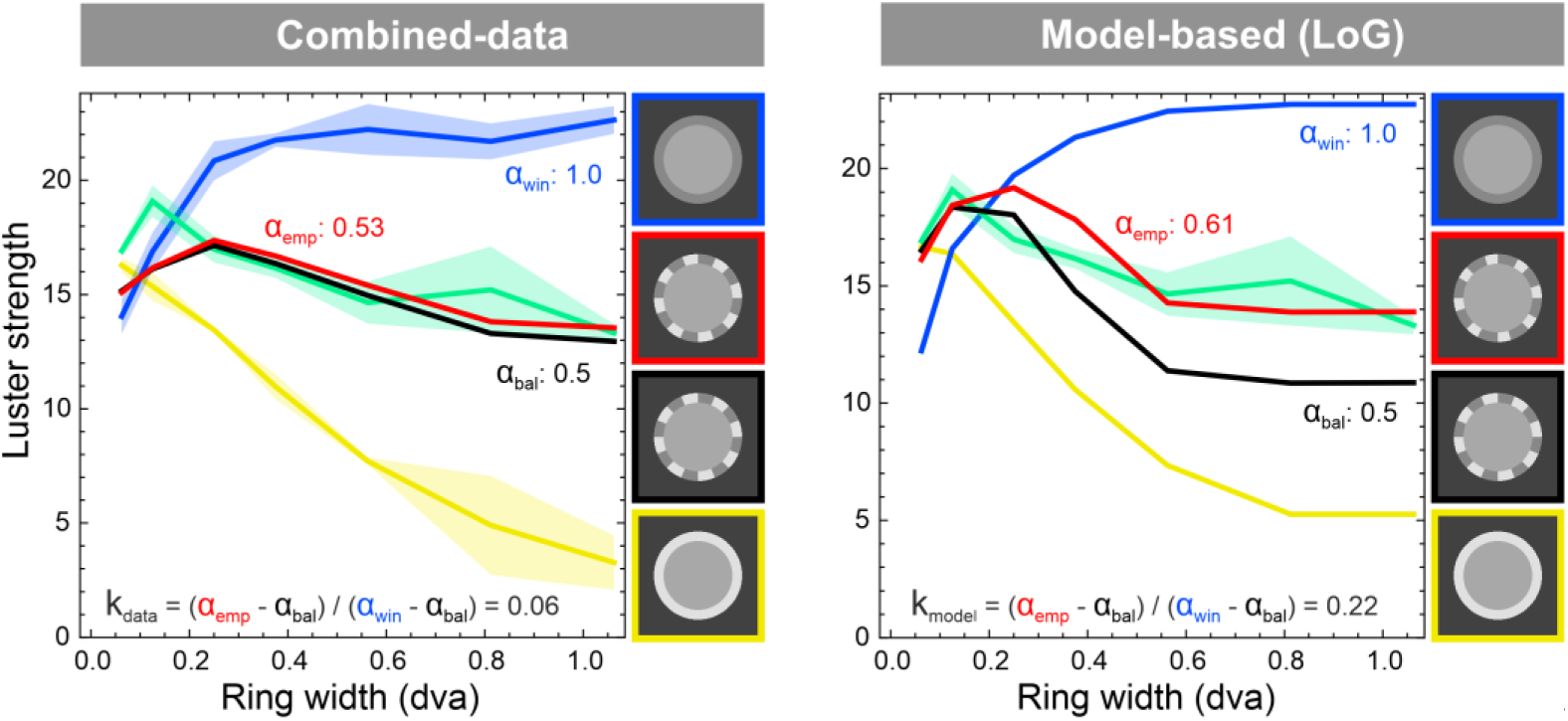
Illustration of how the indices *k_data_* (left) and *k_model_* (right) are calculated. As an example, we use the luster curve from subject S01 of the first stimulus set where the mixed-ring condition had 16 segments and a spatial proportion of *α_ring_* = 0.5 (stimulus with black frame). For the combined-data method, the empirical luster curve (green) was fitted with a convex combination of the two curves that represent the pure effects (yellow for the weaker effect, blue for the stronger effect). For the model-based method, the parameter *α_emp_* represents the spatial proportion of that specific stimulus (with varying ring widths), for which the model calculates a curve (red) that best fits the empirical luster curve (green). In both cases, index *k* represents how strongly the empirical luster curve deviates from the curve representing the prediction for the balanced integration method (*α_bal_*, black), relative to the prediction for the winner-takes-all integration method (*α_win_*, blue, which always has a value of 1.0): *k =* (*α_emp-_ α_bal_*) / (1.0 – *α_bal_*).

For the model-based methods, we had to a use a different approach to determine the best fit for the empirical luster curve, as this method requires actual stimuli to calculate a model output see (Appendix A). We therefore needed additional stimuli in addition to those used in the experiment: For each segment number and each ring width, we generated mixed-ring stimuli with spatial proportions *α_i_* for the two ring parts between 0.0 and 1.0 in steps of 0.01 leading in each case to 101 candidate curves. We then determined for each empirical luster curve, which of the 101 curves best fits the empirical curve and set *α_emp_* to the corresponding *α_i_* and calculated the index *k_model_* in the same way as *k_data_* (see the right diagram in Fig. 8).

Figure 9 shows the different types of *k* indices (columns) separately for the two groups **A+** (top row) and **A-** (bottom row). For each combination of the four different segment numbers (differently colored lines) and the three different ring proportions (abscissa), the data points represent the average *k* index across the three subjects of a group and the two stimulus sets, as these two sets produced very similar trends in the luster data (see Appendix D). A first look at the data confirms our previous observations: The stronger contrast effect in the mixed-ring stimuli contributes more to the overall luster impression than the weaker effect, as all *k* values are above the zero line, which represents a balanced integration process. However, the weight of the stronger effect differs significantly between the two groups of subjects. In group **A-**, the weights are generally much higher than in group **A+**. This is particularly evident in the *k* indices calculated using the model-based method. For both the LoG kernel (middle column) and the Gabor kernel (right column), the *k_model_* values in group **A+** are always less than 0.3 and in group **A-** always greater than 0.8. Generally, this trend can also be observed in the *k* indices calculated with the combined-data method (first column). However, especially for group **A-**, the range of *k_data_* values is considerably larger than that of the *k_model_* values. At least in group **A-**, the *k_data_* values also show a clear dependence on the number of ring segments—an effect that is much less pronounced in the corresponding *k_model_* indices. Specifically, increasing the number of ring segments leads to lower *k_data_* indices, indicating that the weights of the stronger and the weaker effect become more balanced as the number of segments in the mixed-ring stimuli increases. We will provide two alternative explanations for this effect in the discussion section (2.6.).

**Figure 9:**
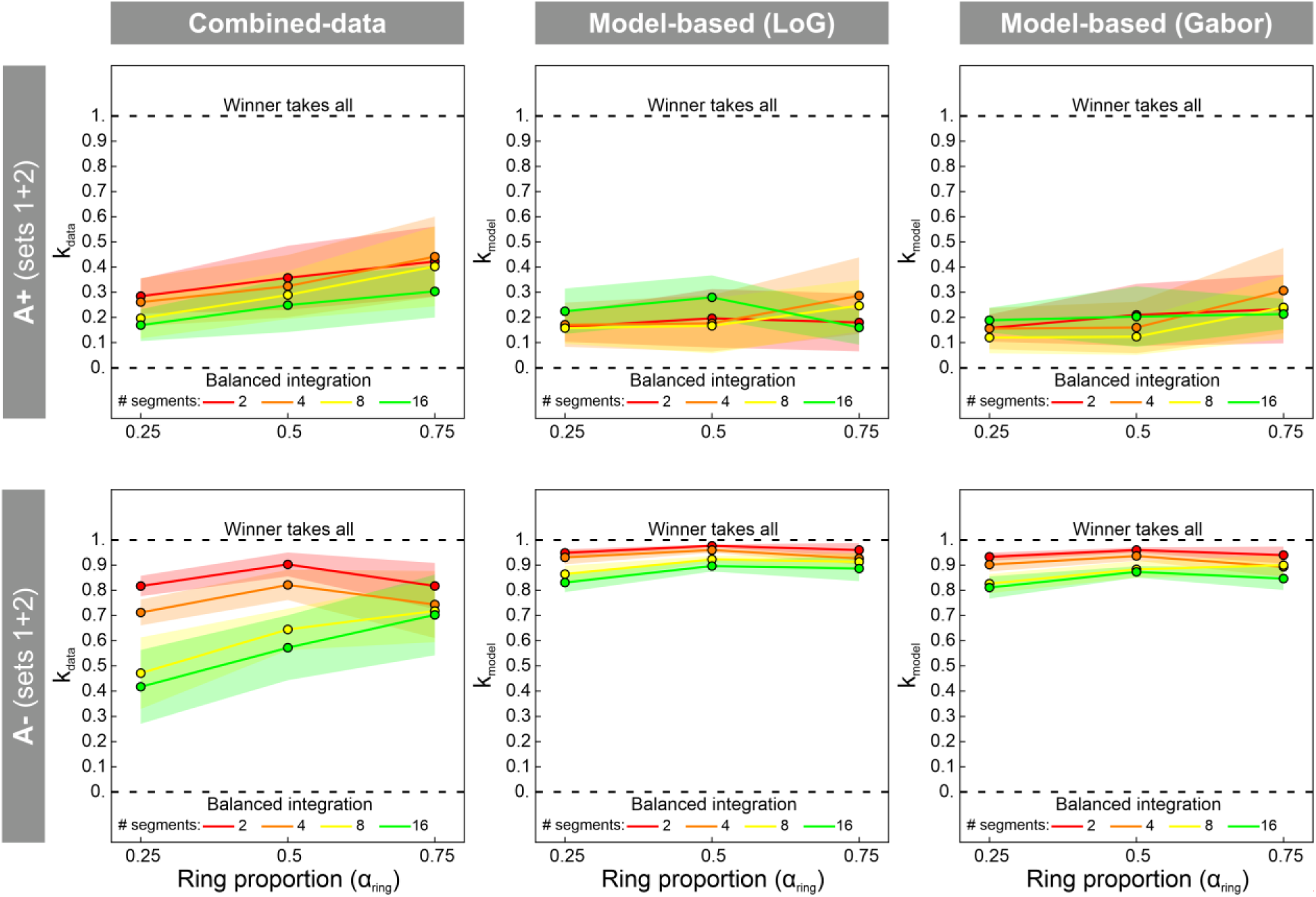
The different types of *k* indices calculated using either the combined-data method (left column) or the model-based method (middle column for the LoG, right column for the oriented ON-OFF model), separately for the two subject groups **A+** (top row) and **A-** (bottom row). In the diagrams, the data points represent the *k* values for each combination of the three different ring proportions (abscissa) and the four different segment numbers (colored lines), averaged across the three subjects in a group and the two stimulus sets. Transparent areas represent the SEM in both directions.

Figure 10 shows the results of the homogeneity rating task, separately for the two groups **A+** and **A-** (note that homogeneity ratings were only obtained for those stimuli that were classified as lustrous during the first part of each trial, see Section 2.3). For the statistical testing of the rating data, we performed a three-way ANOVA on the combined datasets that include both groups with the factors group (**A+** and **A-**), number of segments (1, 2, 4, 8, and 16) and ring proportion (0.0, 0.25, 0.5, 0.75, and 1.0), which revealed significant effects for all three main effects and all three first-order interaction effects (see Table E1 in Appendix E). In addition, we performed a two-way ANOVA separately for groups **A+** and **A-** with the factors number of segments and ring proportion. As we were particularly interested in how these factors interact with regard to the mixed-ring conditions, we tested the ring proportion variable only with the levels 0.25, 0.5, and 0.75 and the segment number variable with the levels 2, 4, 8, and 16. Again, all main effects and interaction effects were significant (see Table E1).

**Figure 10:**
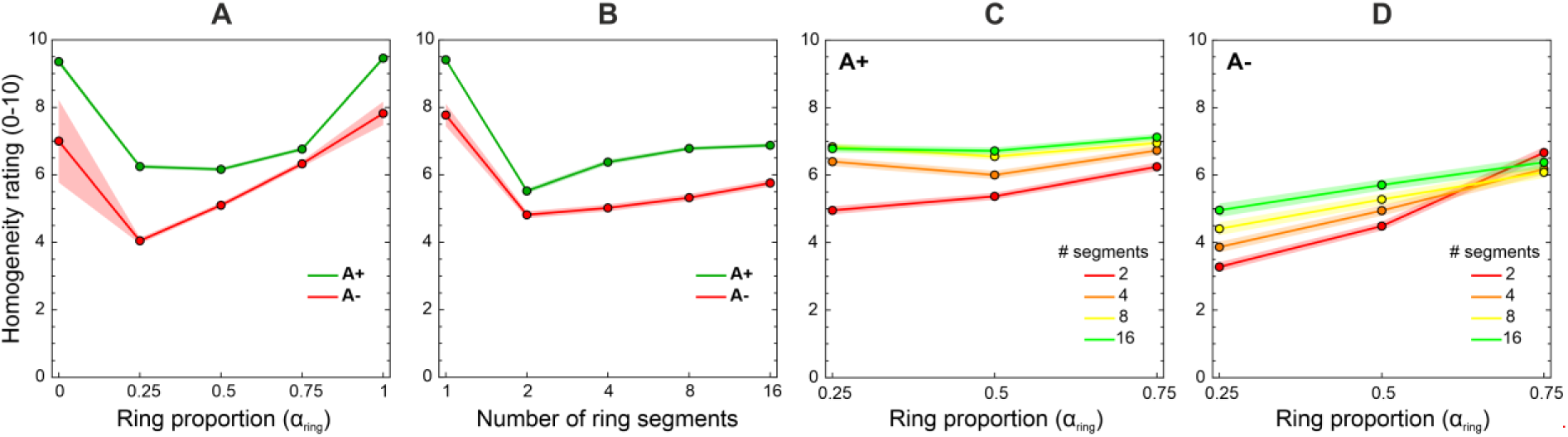
Results of the homogeneity rating task. Diagrams A and B show the mean homogeneity ratings depending on the spatial proportion between the two ring parts (A) and the number of ring segments (B), separately for the two groups **A+** (green lines) and **A-** (red lines). Diagrams C and D show the interactions between these two stimulus parameters exclusively for the mixed-ring conditions for group **A+** (diagram C) and group **A-** (diagram D). The transparent areas represent the SEM in both directions.

Panels A and B in Figure 10 reveal two very clear trends. First, both groups perceived the luster in the target area of the mixed-ring stimuli (with spatial proportions between 0.25 and 0.75 or segment numbers greater than 1, respectively) as significantly less homogeneous than in the two pure conditions with a full ring. Second, spatial homogeneity of the lustrous impression was generally rated significantly lower by group **A-** than by group **A+**. This difference is evident even for the two pure conditions, for which spatial luster homogeneity was rated at an average of only 7.77, whereas group **A+** rated these stimuli as almost perfectly homogeneous, with an average rating of 9.41. When considering only the mixed-ring stimuli (Panels C and D in Fig. 10), a general dependence on the number of ring segments emerges, similar to that observed for the *k* indices (see Fig. 9): The greater the number of ring segments, the more homogeneous the luster in the target area appears. In addition, at least for group **A-**, perceived homogeneity appears to increase systematically with increasing ring proportion (Fig. 10 D).

### 2.6. Discussion

The results of the matching task suggest that the two subject groups **A+** and **A-** differ significantly in how they spatially integrate different contrast effects on binocular luster. Neither a strict winner-takes-all integration nor a pure balanced integration, which assumes equal weights for the two different contrast effects, can explain these results.

To test the assumption of a balanced integration, we used different methods to calculate corresponding luster predictions for the mixed-ring stimuli (see Appendix D), namely a combined-data method and two model-based methods, which are based on the LoG ON-OFF model and the oriented ON-OFF model variant, respectively (see section 2.4.). None of these models was able to accurately predict the empirical luster settings. The best results, in terms of the coefficient of determination R^2^, were obtained for group **A+**, which shows a much stronger tendency toward balanced integration than group **A-** (see Fig. 9): The luster settings for the mixed-ring conditions of group **A+** could best be predicted by the oriented ON-OFF model (with R^2^ = 0.31), while the predictions using the LoG ON-OFF model (R^2^ < 0) and the combined-data method (R^2^ = 0.01) were significantly worse. For the mixed-ring data of group **A-**, all of these methods yielded an R^2^ < 0. Note that R^2^ values can be negative if the mean of the dataset (in our case, the luster settings) provides a better prediction than the model, i.e., if the sum of squared errors (SSE) is greater than the total sum of squares (SST).

Our data therefore suggest an unbalanced integration of the two contrast effects, where both effects contribute to the lustrous impression, but with different weights. In both groups, the stronger of the two contrast effects has a higher weight than the weaker one; however, this weighting asymmetry is considerably more pronounced for group **A-** than for **A+**. This trend is consistent across the different analysis methods employed.

We next investigate why the *k_data_* and the *k_model_* indices differ. By construction, the combined-data method produces luster curves that are equally spaced and do not depend on the number of ring segments (see the lines in the top panels of Figs. D1 and D2 in Appendix D). In contrast, the model-based predictions show shifts that are not only influenced by the number of ring segments but also clearly differ between groups **A+** and **A-**. They also differ to some extent between the two variants of the model, especially at higher ring segment numbers (compare the corresponding lines between the middle and the bottom panels of Figs. D1 and D2). These shifts are caused by two different components of the model. To illustrate the respective effects in more detail, Figure 11 shows the predictions calculated with the LoG ON-OFF model for stimuli of set 2 using eleven different ring proportions between 0 and 1 in steps of 0.1, separately for the two groups **A+** (top row) and **A-** (bottom row).

**Figure 11:**
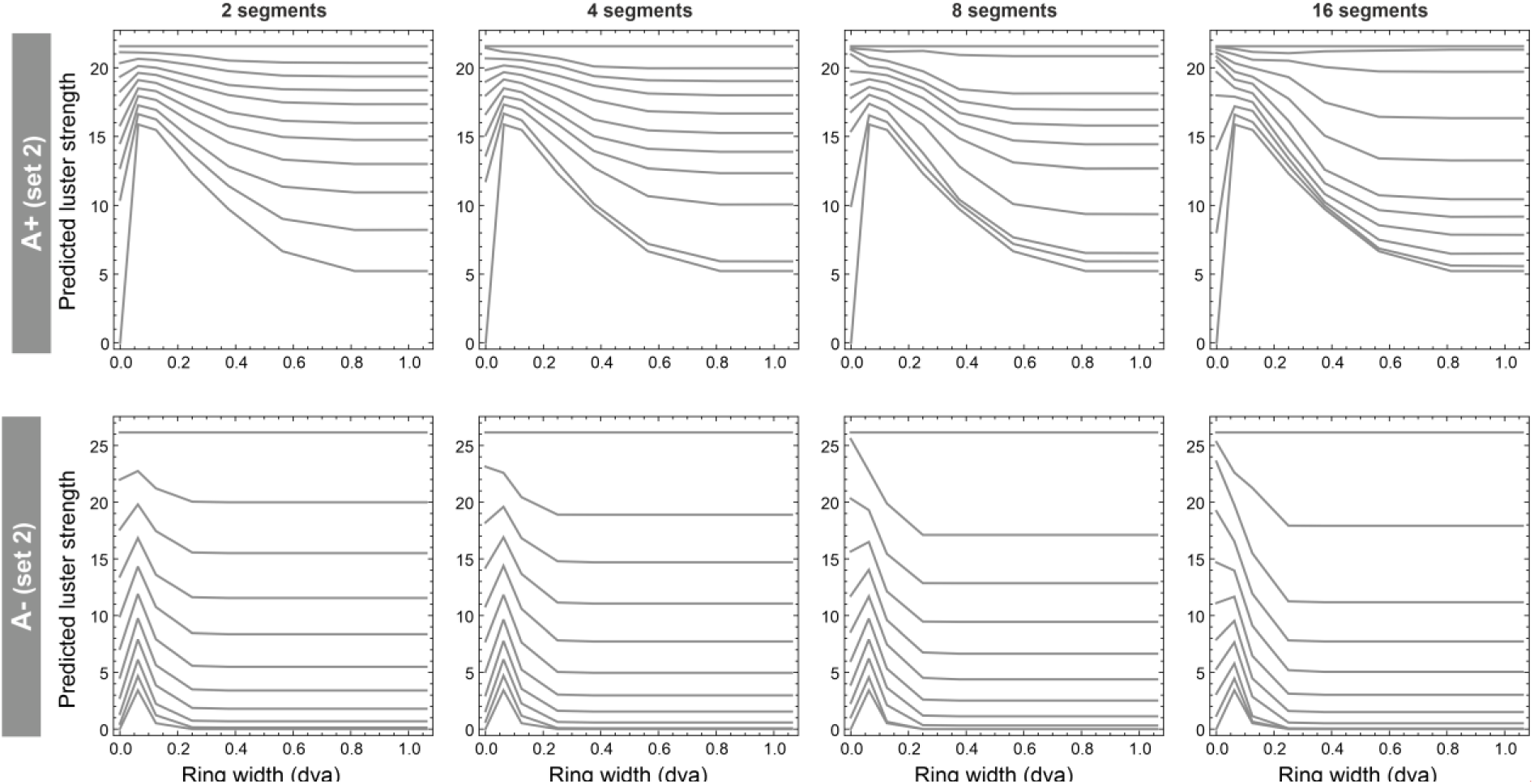
Curve shifts in the model-based predictions. The diagrams show the predicted luster curves for stimuli of set 2 based on the LoG ON-OFF model for eleven different ring proportions (between 0.0 and 1.0 in steps of 0.1) and the four different segment numbers (columns), separately for group **A+** (top row) and **A-** (bottom row). The higher the ring proportion, the higher the magnitude of the model prediction. However, the increase in this magnitude is not proportional to the increase in ring proportion but depends on the different transducer functions of the groups and is also affected by the number of ring segments.

The first effect is most clearly observed in the diagrams showing the prediction curves for stimuli with only two ring segments, because these curves are only weakly affected by the second effect, which depends on the number of ring segments (left column in Fig. 11). For group **A+**, the curves become denser with increasing ring proportion (that is, at higher luster strengths), whereas for group **A-**, the curves become denser with decreasing ring proportion (at lower luster strengths). These effects are caused by the different transducer functions of the two groups (see Fig. 4). Due to the decelerating transducer of group **A+**, weaker interocular conflicts are weighted higher compared to stronger conflicts, whereas for group **A-**, the accelerating transducer causes the opposite effect. This implies that the transducers reflect group-specific sensitivities to interocular discrepancies. Group **A+** is much more sensitive to weaker than to stronger interocular conflicts, whereas the reverse holds for group **A-**.

The second effect underlying the shifts of the luster curves strongly depends on the number of ring segments and has its origin at the level of the monocular contrast detectors, which in our model are simulated either with an LoG or a Garbor filter kernel. As shown in Figure 11, with increasing number of ring segments the luster curves tend to move downward. That is, the more ring segments, the weaker the lustrous impression becomes—an effect that has the same direction for both subject groups. As mentioned in the model description (Appendix A), the magnitude of the lustrous response mainly depends on the relative proportion of local conflicts produced by ON-OFF pairings, whereas ON-ON and OFF-OFF pairings contribute much less to the overall response (Fig. A2 in Appendix A). Figure 12 illustrates how the proportion of these ON-OFF pairings depends on the number of ring segments, using example stimuli from stimulus set 1. Figure 12A illustrates which types of monocular contrast mechanisms are activated at different locations within the target area of the stimulus. OFF-center mechanisms are stimulated at corresponding positions when the filter kernels exclusively cover areas that include the lighter ring part (associated with the weaker *Consistent A* condition, see the red areas in Fig. 12). Stronger ON-OFF signals are generated at those locations, at which the filter kernels cover areas that only contain the darker ring part (associated with the *Reversed B* condition, see the yellow areas in Fig. 12). The critical stimulus locations responsible for the reduction of ON-OFF signals are intersections between the two different ring parts where two different ring luminances are spatially integrated by the filter kernels. In these cases, slightly less ON-OFF pairings than pairings with equal polarities are produced. For the stimulus example with two ring segments shown in Figure 12A, the proportion of ON-OFF pairings is about 48%. With increasing number of ring segments, and therefore with increasing number of such intersections, the relative number of ON-OFF pairings continuously decreases until it reaches a proportion of less than 36% with rings having 16 segments (Fig. 12B). Note that this effect also depends on the size of the filter kernel. At some locations within the target area, larger kernels can even cover more than two ring segments. This is especially the case for group **A+**, whose estimated model parameters include comparatively large kernel radii. The fact that, for both groups, the sizes of the kernels are always much larger for the Gabor than for the LoG filter (see Fig. 4), also explains the differences between corresponding prediction curves, particularly for stimuli with more than four ring segments (see Figs. D1 and D2 in Appendix D).

**Figure 12:**
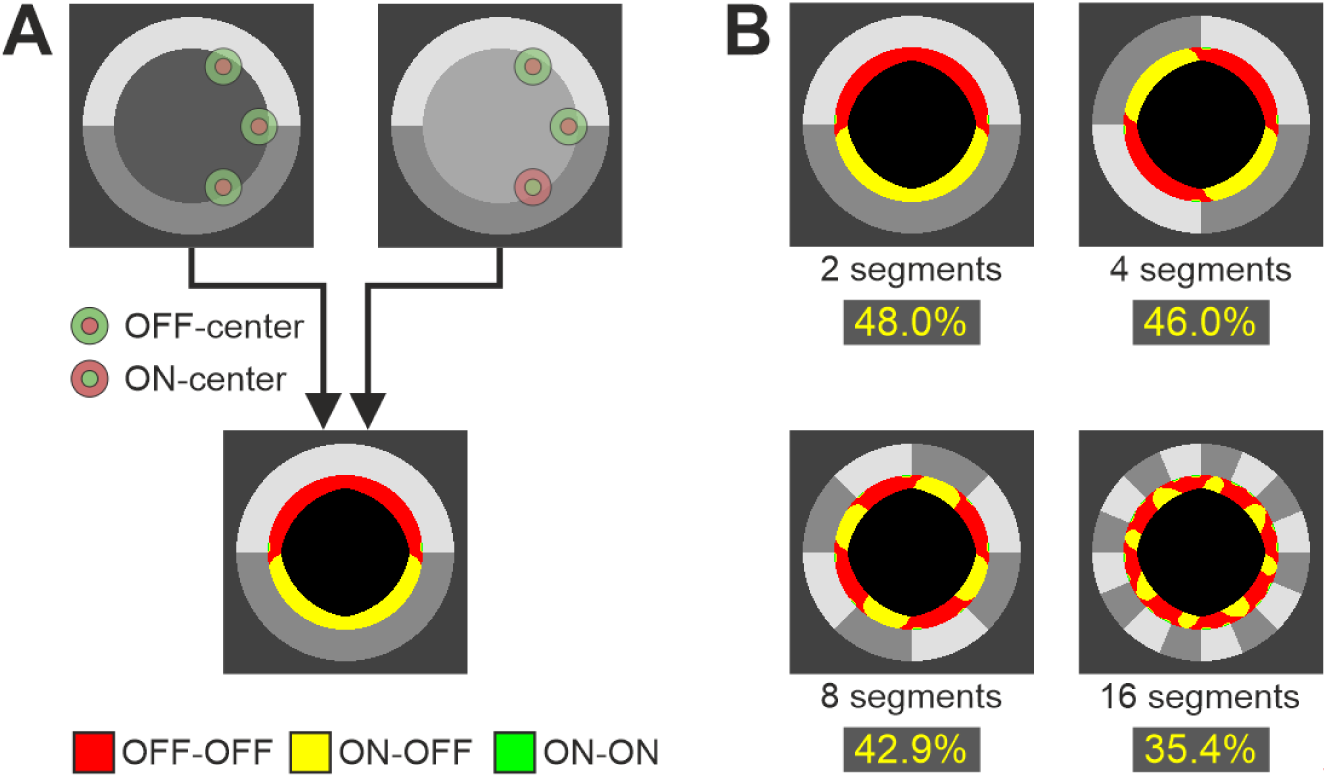
Why do the model-based predictions depend on the number of ring segments? (A) Example stimulus pair from the first stimulus set with a ring subdivided into two segments. Filter kernels that exclusively cover one ring part either produce weak OFF-OFF pairings (red areas) or strong ON-OFF pairings (yellow areas). At the intersections between two ring parts, at which two different ring luminances are spatially integrated by the filter kernel, more OFF-OFF than ON-OFF pairings are produced. (B) The proportion of ON-OFF signals continuously decreases with increasing number of ring segments, or number of intersections, respectively. Since the magnitude of the model output mainly depends on the number of strong ON-OFF signals, this explains why the model predicts weaker lustrous responses for stimuli with increasing number of ring segments.

The differences between the *k_data_* and the *k_model_* indices are therefore due to the fact the model-based method predicts curve shifts that depend on both the number of ring segments and the different sensitivities to the luster phenomenon, whereas the combined-data method does not take either of these influences into account. At least with regard to the number of ring segments, the empirical data indicate that the magnitude of the lustrous response does indeed depend on this variable. In particular, the results of group **A-** clearly show that the lustrous impression becomes increasingly weaker with increasing number of ring segments—an effect that was correctly predicted by the model.

However, an alternative interpretation for this effect exists that is not based on the properties of our model. The results of the homogeneity task suggest that the number of ring segments also has an influence on the spatial homogeneity of the luster (Fig. 10). As the number of segments increases, the luster within the target area is increasingly perceived as spatially uniform. This could mean that in mixed-ring stimuli with only a few segments, the observers can clearly distinguish between areas with different degrees of luster. As we have already speculated in the Methods section (see 2.3. Procedure), the observers may then (consciously or unconsciously) base their judgments more strongly on those areas with stronger luster. As the number of ring segments increases, the two effects may gradually merge into a spatially more homogeneous luster impression, in which the weighting of the two effects becomes more balanced. The stronger impact of the number of ring segments observed in group **A-**could be due to the fact that the perceived difference between the two effects is much greater for group **A-** than for group **A+**. In group **A-**, the stronger effect (caused by those ring parts associated with the *Reversed B* condition) produces a very strong lustrous response, while the weaker effect (associated with the *Consistent A* condition) hardly evokes any impression of luster. In contrast, in group **A+**, both effects generally show some degree of luster.

Interestingly, though, the subjects reported that the homogeneity task was the most difficult and the most time-consuming part of a trial. While they had no problems matching the strength of the lustrous impressions, they described the evaluation of the spatial homogeneity of the luster as a very artificial and abstract task that required direct comparisons between different areas of the stimulus. This could indicate that a greater weighting of the stronger effect during the matching task was, at least, not a conscious strategy.

## 3. General Discussion

The aim of this study was to investigate how the visual system processes complex luster stimuli in which two different contrast effects are combined. For this purpose, we used dichoptic center-ring-surround stimuli with a ring element that was divided into a varying number of segments, where neighboring ring parts differed in their luminance and in the strength of the effect on perceived luster. We tested in particular how the two different contrast effects are spatially integrated by the visual system.

The results from both the matching experiment and the lustrous/non-lustrous classification task, revealed significant individual differences in sensitivity to binocular luster. We identified two main trends in the data, where one group of subjects (**A-**) strongly responded to stimuli from the pure *Reversed B* contrast condition but barely to stimuli from the other condition *Consistent A*, whereas the other group (**A+**) strongly responded to stimuli from both conditions. For both groups, the fit of our interocular conflict model to the luster data was excellent for each of the two basic contrast conditions. However, the parameter values of the best fit differed significantly between the two groups, in particular with regard to the transducer functions, which had a decelerating shape for group **A+** and an accelerating shape for group **A-**. These results indicate that subjects from group **A+** can better discriminate low degrees of interocular conflict, whereas subjects from group **A-** are much more sensitive to differences at higher degrees of interocular conflict (see also Fig. 11). In our recent studies on binocular luster with different groups of participants, we only found transducer functions comparable to those of group **A+** (Wendt & Faul, 2022a, 2025). They were also compatible with results from Kingdom and colleagues (2019) who measured dichoptic contrast difference thresholds. However, this is actually not the first time that we observed different individual sensitivities to luster. In an earlier study (Wendt & Faul, 2019), we directly determined psychophysical luster scales using simple dichoptic center-surround stimuli in an inc-dec arrangement where the luminance difference between corresponding center patches was systematically varied. The resulting scaling functions, which were fitted with power functions, showed large differences across subjects with exponents between 0.187 (that is, with a strongly decelerating shape) and 3.136 (strongly accelerating function).

In the present study, the two groups **A+** and **A-** also differed significantly in their luster settings for the mixed-ring stimuli. Our analyses of these data suggest that in both groups, the two different contrast effects are integrated by an unbalanced integration, in which the stronger effect contributes more to the overall lustrous impression than the weaker effect. However, the weight of the stronger effect was much greater for group **A-** than for group **A+**.

Regarding the performance of the ON-OFF model, it turned out that, although the luster data obtained with the two pure contrast conditions could be accurately predicted (see Fig. 4 and Fig. C1), it failed to predict the lustrous impressions elicited by the mixed-ring stimuli. This is simply due to the fact that the model assumes equal weights for different contrast effects, while the empirical data suggest an unbalanced integration process of the underlying mechanism.

To accurately predict the lustrous appearance of the mixed-ring stimuli, the model would first require an additional parameter that represents the imbalance between the weights for the two different contrast effects—which is exactly what the *k*-index does (see Fig. 9). As we have seen in Appendix A, the basic elements of our model are local binocular conflict signals (s. step 2 in Fig. A1). These local conflict signals are the units to which the different weights for the different contrast effects would have to be assigned. However, there is a problem: If we consider an individual binocular conflict signal at a given location within the target area, what weighting does it receive? In our present study, in which the two ring parts generally also produce different types of interocular polarity pairings, with the stronger contrast effect associated with ON-OFF pairings and the weaker effect associated with OFF-OFF pairings (see Fig. 12), the answer seems simple: Assign the stronger weight to those conflict signals that are produced by monocular ON-OFF pairings and the weaker weight to those conflict signals that result from OFF-OFF pairings. However, as we have shown in Figure A2, ON-OFF signals are not produced exclusively by the ring luminance that represents the stronger contrast effect, but also, depending on the width of the ring, by the other ring luminance, so this would be an inappropriate procedure. Furthermore, if we would use two different ring parts that differ in luminance but produce the same type of polarity pairing, this would also fail. In other words, the local conflict signal alone does not reveal whether it was generated by a stronger or a weaker contrast effect. This means that more information, based on comparisons between local conflict signals across the entire set of signals, would be required to determine the weight for an individual conflict signal. Constructing a corresponding model component would therefore be a major challenge and it is questionable whether such a mechanism has a physiological counterpart at a low level of visual processing.

It is therefore likely that higher-level processes are involved here: At least in the case of stimuli with a spatially uniform target area, as those used in most of our studies, binocular luster is assumed to be a filling-in phenomenon originating at the contour of the target area and then spreading to the remaining parts (Wendt & Faul, 2020; Zöller, 1998; for a different type of luster stimuli in which the target area comprises a complex pattern in the form of horizontal luminance gratings, see Georgeson et al., 2016; Kingdom, Mohammad-Ali, Breuil, Chang-Ou, & Irgaliyev, 2023; Kingdom et al., 2018; Kingdom et al., 2019). In fact, as can be seen in Figures A1 and A2 in Appendix A, local conflict signals only occur in a narrow strip along the luminance edge of the target patch; the lustrous quality, on the other hand, extends across the entire area of the patch. It is therefore not unreasonable to assume that the spatial integration of various contrast effects takes place at the same level as the filling-in mechanism, i.e., at a level that goes beyond the scope of our model, which serves primarily to explain the early neural processes underlying the phenomenon of binocular luster.

## Acknowledgments

The authors thank Laura Groninger, Angelika Roth, and Theresa Thielemann for their assistance in collecting the experimental data.

Funded by the Deutsche Forschungsgemeinschaft (DFG, German Research Foundation) - 519638685.

## Appendix A The LoG variant of the interocular conflict model of binocular luster (ON-OFF model)

One method to test the assumption that the visual system equally weights both contrast influences in mixed-ring stimuli is to calculate the luster predictions using our interocular conflict model, which implements balanced spatial integration. Since all test stimuli had a target area of constant size, we could use the simple version of the model (Wendt & Faul, 2022a), which comprises a total of five free parameters (a more advanced version that accounts for additional effects due to differences in stimulus size can be found in Wendt & Faul, 2025).

The basic output of the ON-OFF model is a conflict measure *C*′, which represents the magnitude of the lustrous response triggered by a given dichoptic stimulus pair (*I_l_*, *I_r_*) (Fig. A1) and is calculated as follows: First (see step 1 in Fig. A1), the two half-images *I_l_* and *I_r_* of the stimulus pair are convolved with an LoG-kernel that simulates the behavior of monocular contrast detector cells with a circular-symmetric center-surround structure of their receptive fields, the radius of which is one of the parameters fitted with the empirical data using a grid search. The sign of the filter response in the convolved images *I_l_’* and *I_r_’* represents the type of monocular cell that is stimulated by the luminance pattern covered by the cell’s receptive field. A negative sign represents the stimulation of an ON-center cell and a positive sign the stimulation of an OFF-center cell, accordingly. Therefore, each corresponding pair of pixels in the filtered images *I_l_’_(x,y)_* and *I_r_’_(x,y)_* represents one of three possible interocular combinations of ON and OFF signals, i.e. either an ON-ON pairing, an OFF-OFF pairing, or an ON-OFF pairing (see the colored pixels in the binocularly-combined image in Fig. A1).

For further calculations, the model takes only those pixel pairs into account that are located within the target area and whose filter values are both not zero (to avoid rounding errors, the unsigned filter values had to exceed a threshold of 1×10^-6^).

**Figure A1:**
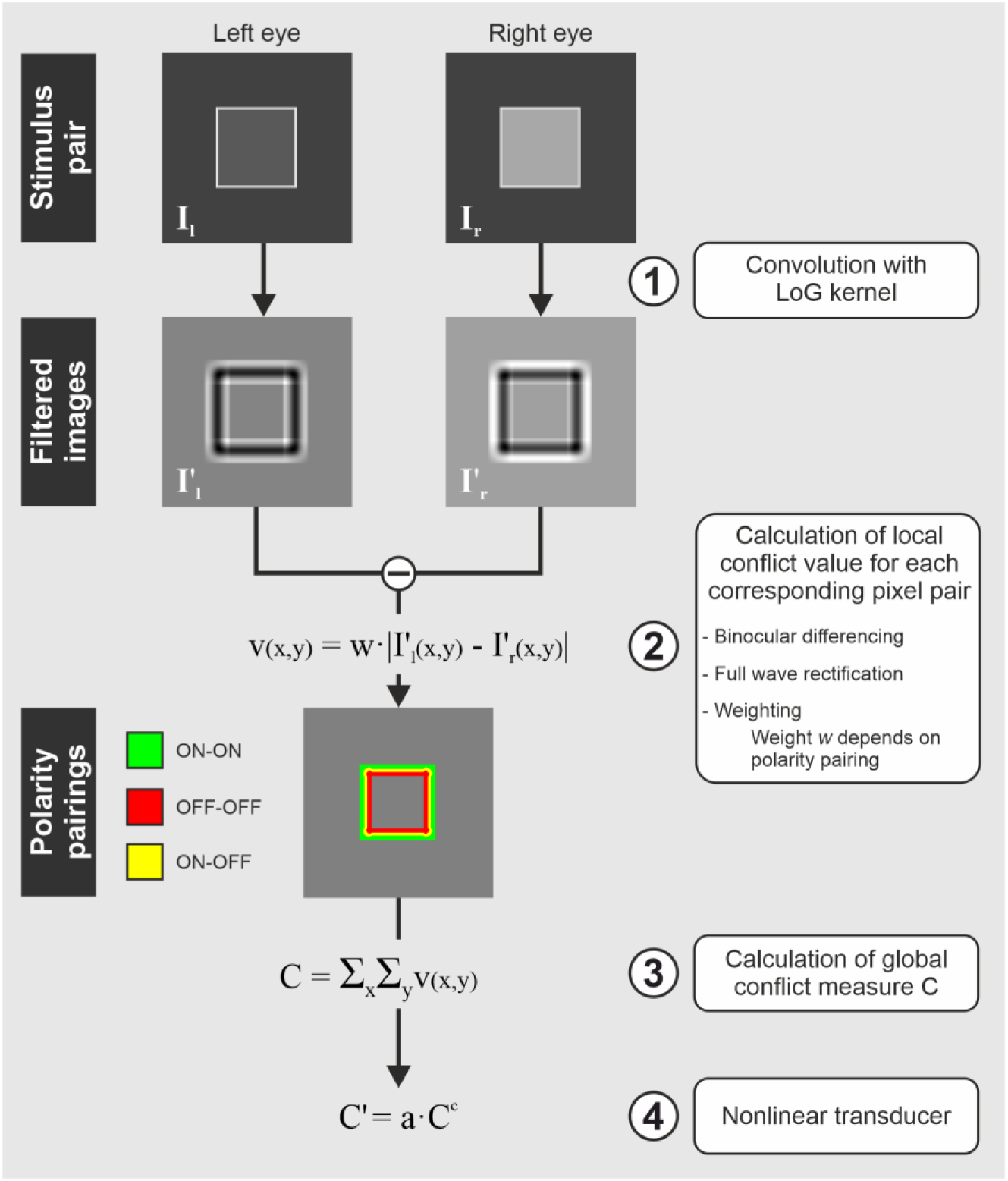
Schematic representation of the LoG variant of the interocular conflict model. (1) The two half-images *I_l_* and *I_r_* of a dichoptic stimulus pair are first convolved with an LoG kernel, simulating the behavior of contrast detector cells with a circular-symmetric center-surround receptive field. Dark pixels in the filtered images *I’_l_* and *I’_r_* represent the stimulation of an ON-center cell, light pixels the stimulation of an OFF-center cell. (2) At the binocular level, corresponding pixel pairs within the target area are then combined into a local binocular conflict signal *v(x,y)*, resulting from the full-wave rectified difference between the two filter responses, weighted by weight *w*, whose magnitude depends on the polarity pairing of the input signals. (3) The local conflict values are then spatially integrated into a global conflict measure *C.* (4) In the final step, a nonlinear transducer is applied to the global conflict measure to produce the adjusted model output *C’*, which represents the magnitude of the lustrous response.

Each of these monocular contrast signal pairings is then combined into a binocular response, which we refer to as the local conflict value *v(x,y)* (step 2 in Fig. A1). Irrespective of the type of polarity pairing, the two signals are integrated by the same binocular mechanism, which is based on a differencing process (Henricksen & Read, 2016; Georgeson, Sato, Chang, & Kingdom, 2025; Kingdom, 2012). In our model, the difference between the two corresponding filter values *I_l_’_(x,y)_* and *I_r_’_(x,y)_* is full-wave rectified and then multiplied with weight *w*. In case the two filter values of a pair had opposite signs (representing the occurrence of an ON-OFF pairing), weight *w_ON-OFF_* was set 1.0. The weights *w_ON-ON_* and *w_OFF-OFF_* for the two remaining cases with equal signs (representing either an ON-ON or an OFF-OFF pairing) are further parameters of our model whose values are determined by a fit with the empirical data. Generally, these two weights are considerably lower than weight *w_ON-OFF_*.

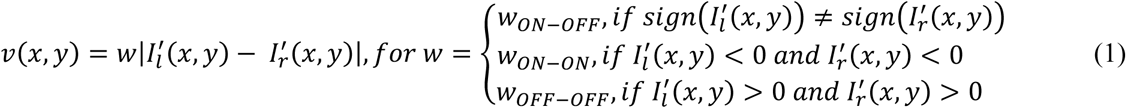

The sum of these local conflict values then represents the global conflict measure *C* (step 3 in Fig. A1):

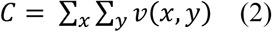

In the final step (see step 4 in Fig. A1), a nonlinear transducer is applied to conflict measure *C* to produce the model output *C*′:

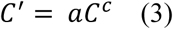

Factor *a* and exponent *c* of the transducer function are the remaining parameters of our model, which are also determined by a fit with the empirical data.

Figure A2 demonstrates how the empirical data of our previous studies with dichoptic center-ring-surround stimuli can be explained by the ON-OFF model, using two luster curves from Figure 1 as examples. For each of the two contrast conditions *Reversed B* (upper panel) and *Consistent A* (lower panel), three different stimulus pairs are shown which differ in the width of the ring element. The third row in each panel shows, how the width of the ring influences the proportions of the three different types of polarity pairings (see the differently colored pixels in the binocularly-combined images). Since the magnitude of the lustrous response is mainly determined by the relative number of ON-OFF pairings, which produce strong conflict signals, this explains the different shapes of the two luster curves. While in the *Reversed B* condition the proportion of ON-OFF pairings increases with increasing ring width, the *Consistent A* condition only produces relevant proportions of these strong signals at lower ring widths.

**Figure A2:**
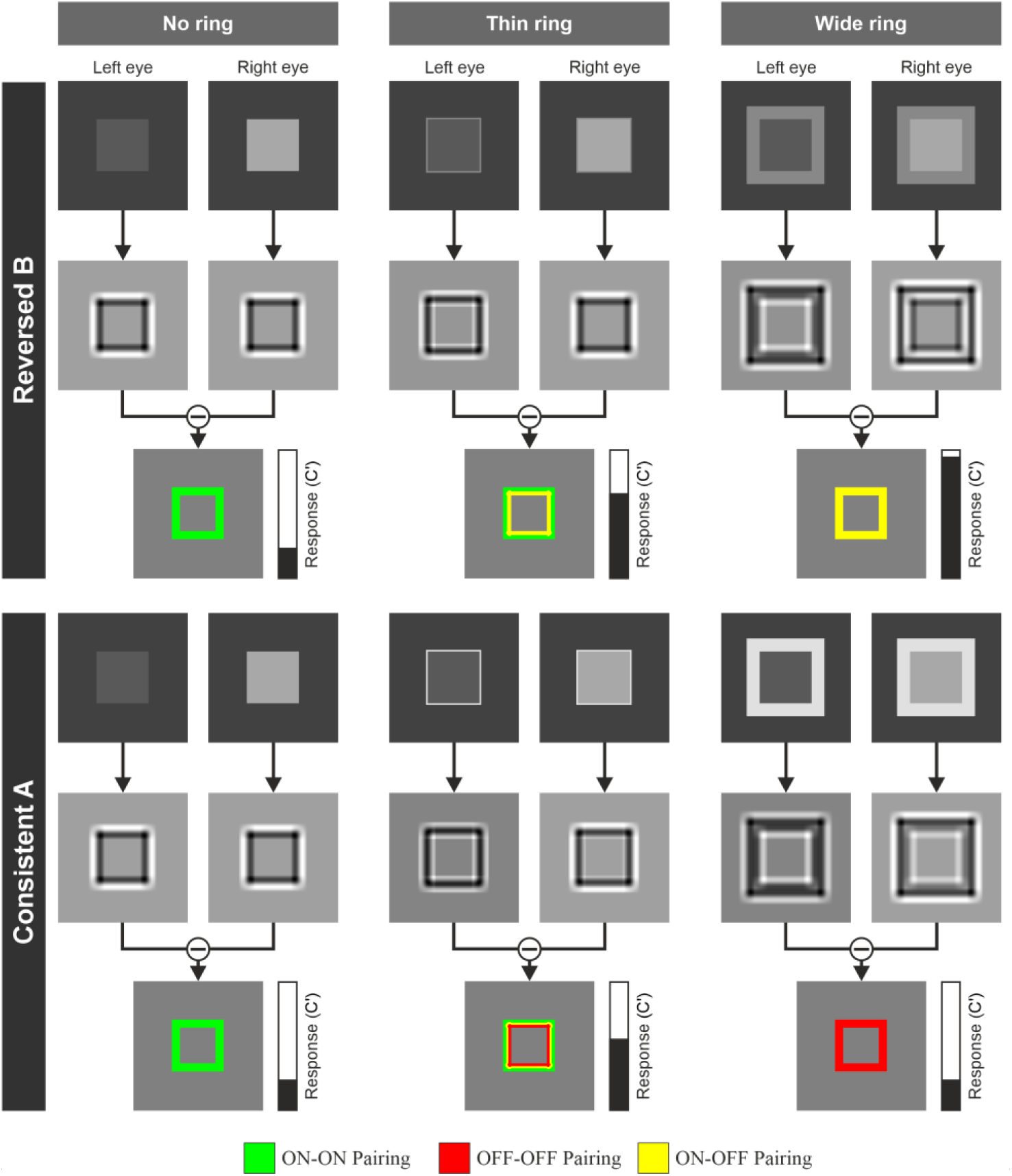
The magnitude of the lustrous response to dichoptic center-ring-surround stimuli can be predicted by the interocular conflict model. **Top half**: Three different stimulus pairs of the *Reversed B* condition (see Fig. 1) are shown, differing in the width of the ring element (top row). As in Figure A1, the middle row shows the filtered images of the two monocular half-images after the convolution with an LoG kernel, which is used to simulate the behavior of contrast detector cells with antagonistic circular-symmetric center-surround receptive fields. Dark pixels represent the stimulation of an ON-center cell, light pixels the stimulation of an OFF-center cell. Pixels at corresponding locations between the two eyes can either produce ON-ON pairings (green pixels in the binocularly combined images in the bottom row), OFF-OFF pairings (red pixels) or ON-OFF pairings (yellow pixels), the latter of which generates much stronger binocular conflict signals than the two other combinations with equal polarities. Since the ring luminance lies between the luminances of the two center patches (see Fig. 1), increasing the ring width increases the amount of strong ON-OFF signals, which in turn leads to a continuous increase in the lustrous response. **Bottom half**: For stimuli of the *Consistent A* contrast condition, where the ring element has a higher luminance than the two center patches, relevant amounts of ON-OFF pairings only occur at smaller ring widths. This explains the non-monotonic shape of the corresponding luster curve (Fig. 1).

## Appendix B The Garbor variant of the interocular conflict model (oriented ON-OFF model)

As a variant of the model, which uses the same five free parameters as the LoG variant (Appendix A), we used Gabor filters in which the antagonistic parts (i.e. filter weights with opposite signs) are spatially arranged side by side. The Gabors had a fixed spatial frequency of 1 cycle/radius. To produce a kernel with an OFF-center structure, we applied a phase shift of π to the Gabor, such that the central part has negative signs and is flanked by parts with positive signs (see Fig. B1-F). In analogy to the LoG-based model, this means that a filter response with a negative sign would represent the stimulation of an ON-center cell and a positive sign the stimulation of an OFF-center cell. For each radius of the Gabor filter, the two half-images of a stimulus pair were convolved with 16 differently oriented kernels (which were sum-normalized to 0 and square-normalized to 1) with orientations between 0° and 168.75° in steps of 11.25° (see the example filter bank in Fig. B1-F).

**Figure B1:**
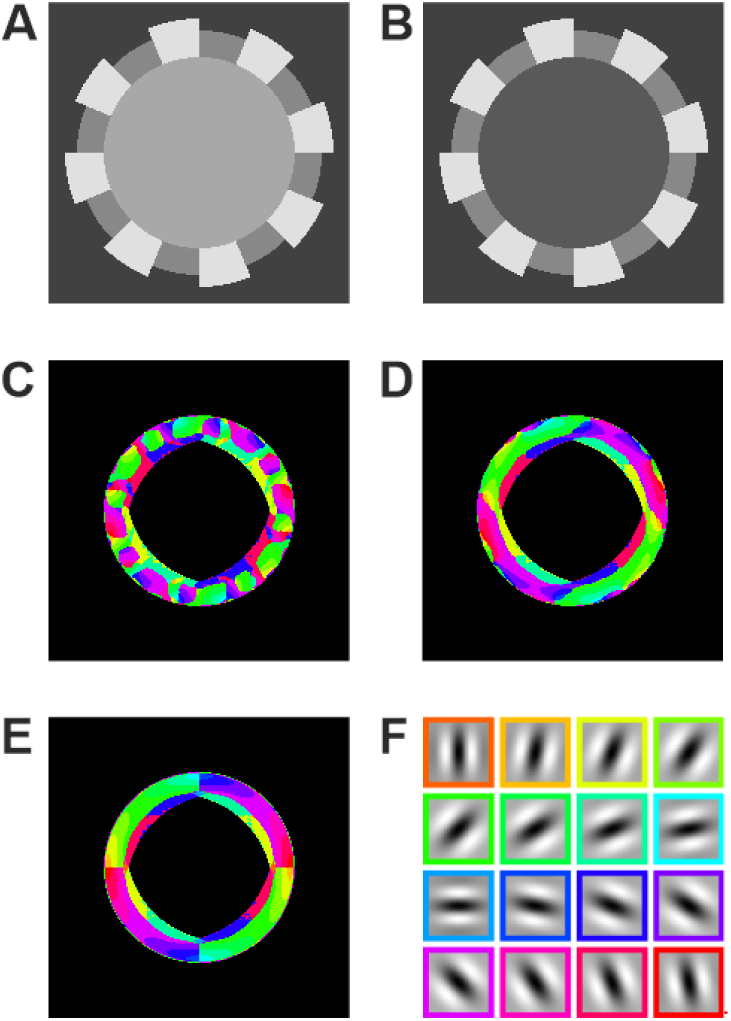
How to combine corresponding monocular signals from Gabor filters? A and B show the left and right half-image of an example dichoptic stimulus pair from stimulus set 2 with 16 ring segments. C and D show, separately for the two eyes, the orientations of those Gabor kernels that produce the strongest absolute signals for each pixel within the central target area (the different colors refer to different orientations of the kernel, see the filter bank in F; the kernel had a radius of 15 pixels in this example). One can see that the Gabor filters with the strongest responses not always have the same orientation at corresponding pixel positions. To calculate local binocular conflict values, our Gabor variant uses the responses from corresponding kernels with identical orientations showing the strongest difference between these signals (as shown in E).

For the binocular level of the Gabor variant, we first had to define a rule how corresponding monocular signals *I’_l,i(x,y)_* and *I’_r,j(x,y)_* (where indices *i* and *j* refer to specific orientations of the kernels) are combined to form a local conflict value *v_(x,y)_*. One variant would be to select the two values from corresponding sets of 16 filter responses that represent the strongest (absolute) filter signals. We investigated whether these two strongest signals are produced by Gabor kernels with same or different orientations and found that, in most cases, the strongest (absolute) signals indeed come from corresponding filters with the same orientation.

However, at many locations within the target area of the stimulus, this was not the case, depending on the size of the Gabor and the complexity of the ring structure of the stimulus (in Fig. B1 we show an example with a more problematic ring structure in this regard). Since we consider it more physiologically plausible that, at the binocular level, monocular cells tuned to the same orientation are paired, we decided to apply a different rule. The local conflict value *v_(x,y)_* therefore results from corresponding signals produced by Gabor kernels with same orientations showing the largest absolute difference between them.

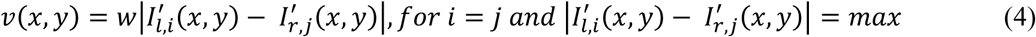

As in the LoG-based model (see Appendix A), weight *w* depends on the interocular contrast polarity combination (*w_ON-OFF_*, *w_ON-ON_* and *w_OFF-OFF_*). The remaining steps for the calculation of the adjusted global conflict measure C’ are identical to those of the LoG model (Appendix A).

## Appendix C Model parameter estimation for the individual data sets

**Figure C1:**
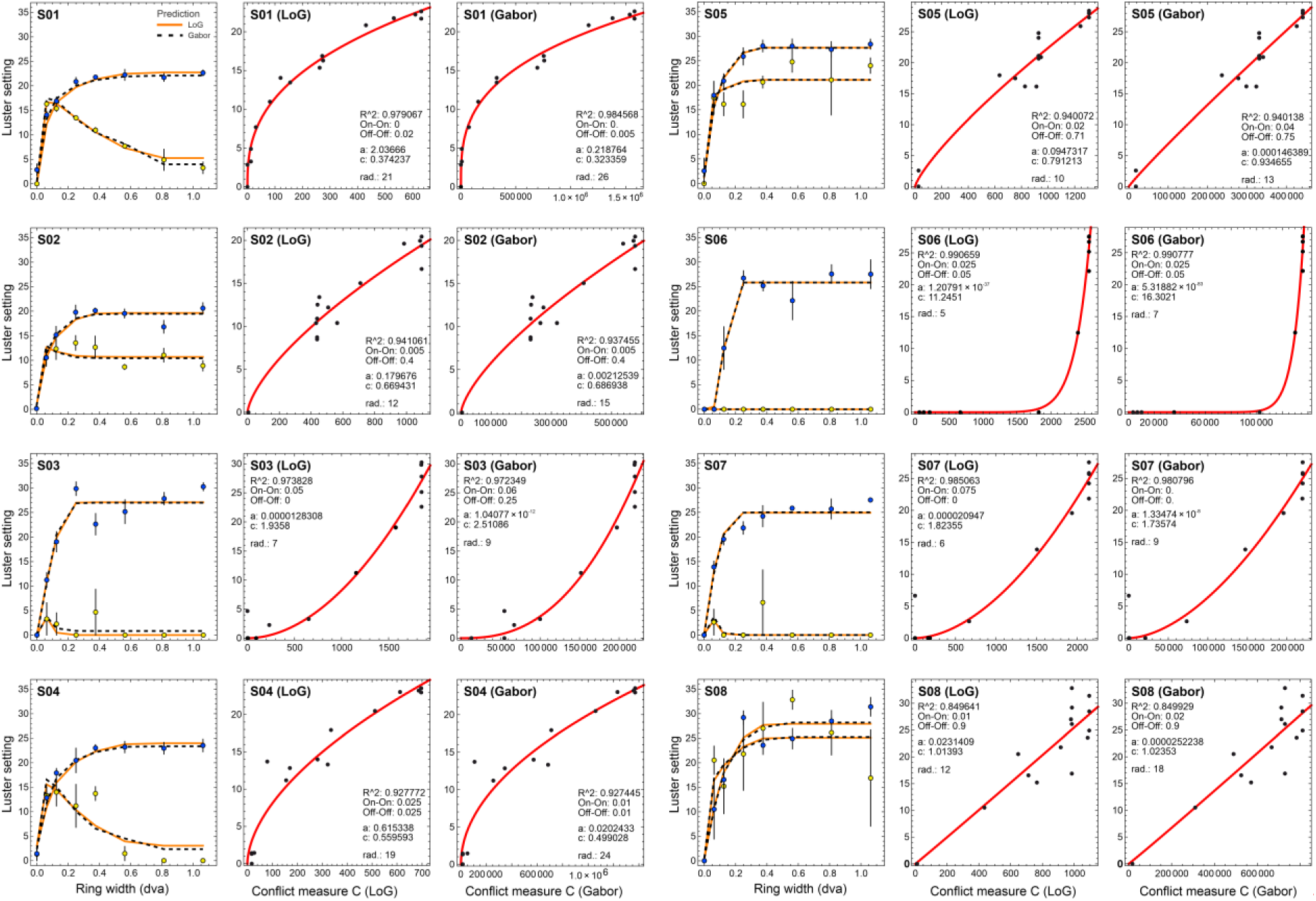
Model parameter estimation for the individual data sets. A set of three diagrams is shown for each of the eight subjects S01 – S08. The left diagram in each set shows the luster settings obtained under the two pure contrast conditions *Consistent A* (yellow symbols) and *Reversed B* (blue symbols) as function of ring width, with the error bars representing the SEM in both directions. In addition, the model predictions based on the best-fitting parameter values are shown for the LoG (orange curves) and the oriented ON-OFF model (dashed curves). The two remaining diagrams of each set show the relationship between the empirical luster settings and the corresponding conflict measures *C* of the LoG variant (middle diagram) and the Gabor variant of the model (right diagram). Note that conflict measure *C* is not the final output of the models (which is *C’*, as it was used for the prediction lines in the left diagrams, s. Appendix A), but an intermediate result which we use here to demonstrate the shape of the transducer functions (red curves).

## Appendix D Predictions for the balanced integration process

**Figure D1:**
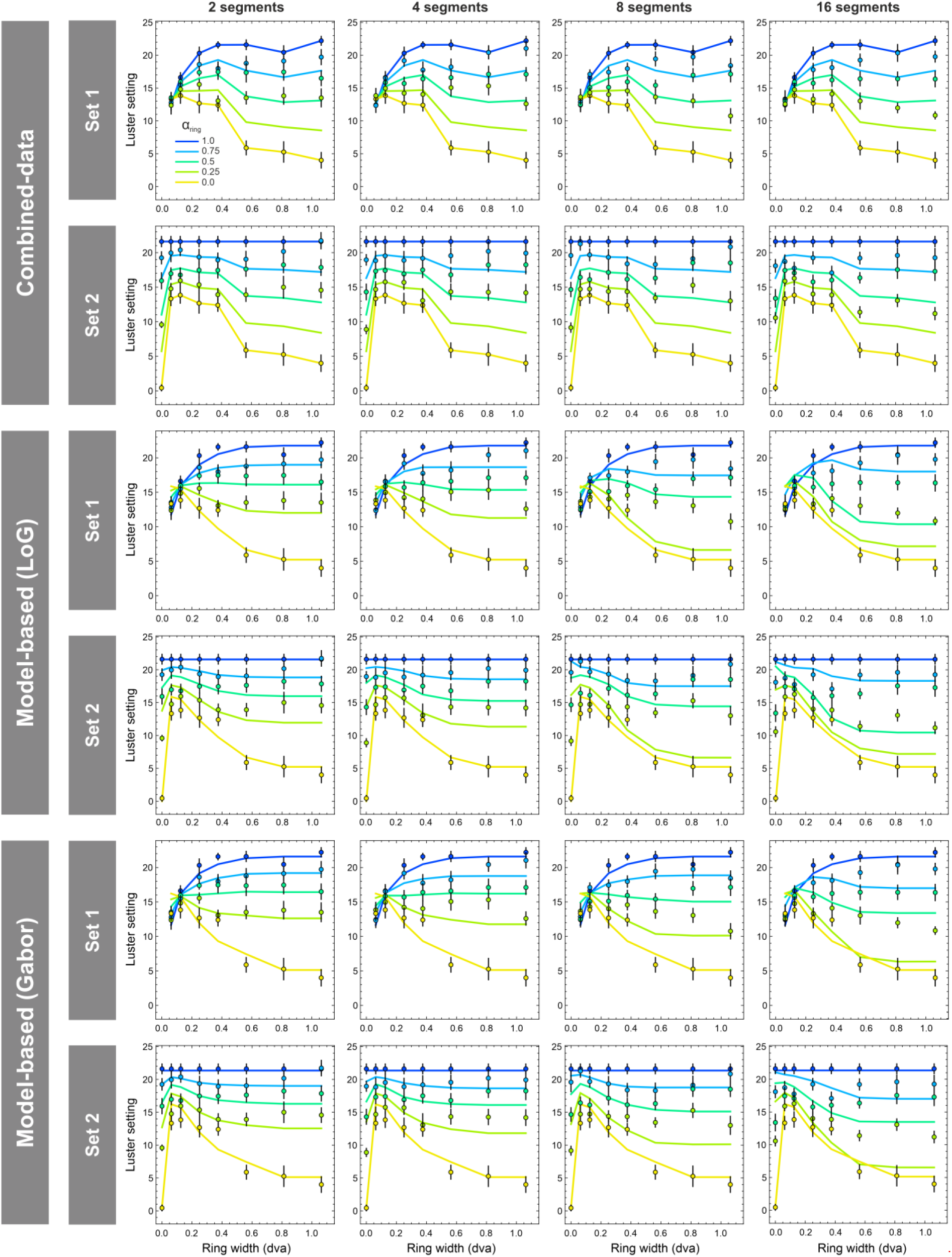
Empirical luster settings (colored disks) and corresponding predictions (colored lines) for group **A+**, separately for the three different methods used to calculate the predictions for the balanced integration process, that is, the combined-data method (top panel), and the two model-based methods, which are either based on an LoG kernel (middle panel) or a Gabor kernel (bottom panel). Each panel shows the results for the two different stimulus sets (rows), separately for the different numbers of ring segments (columns) and the different ring proportions (differently colored symbols and lines). The error bars represent SEM in both directions.

**Figure D2:**
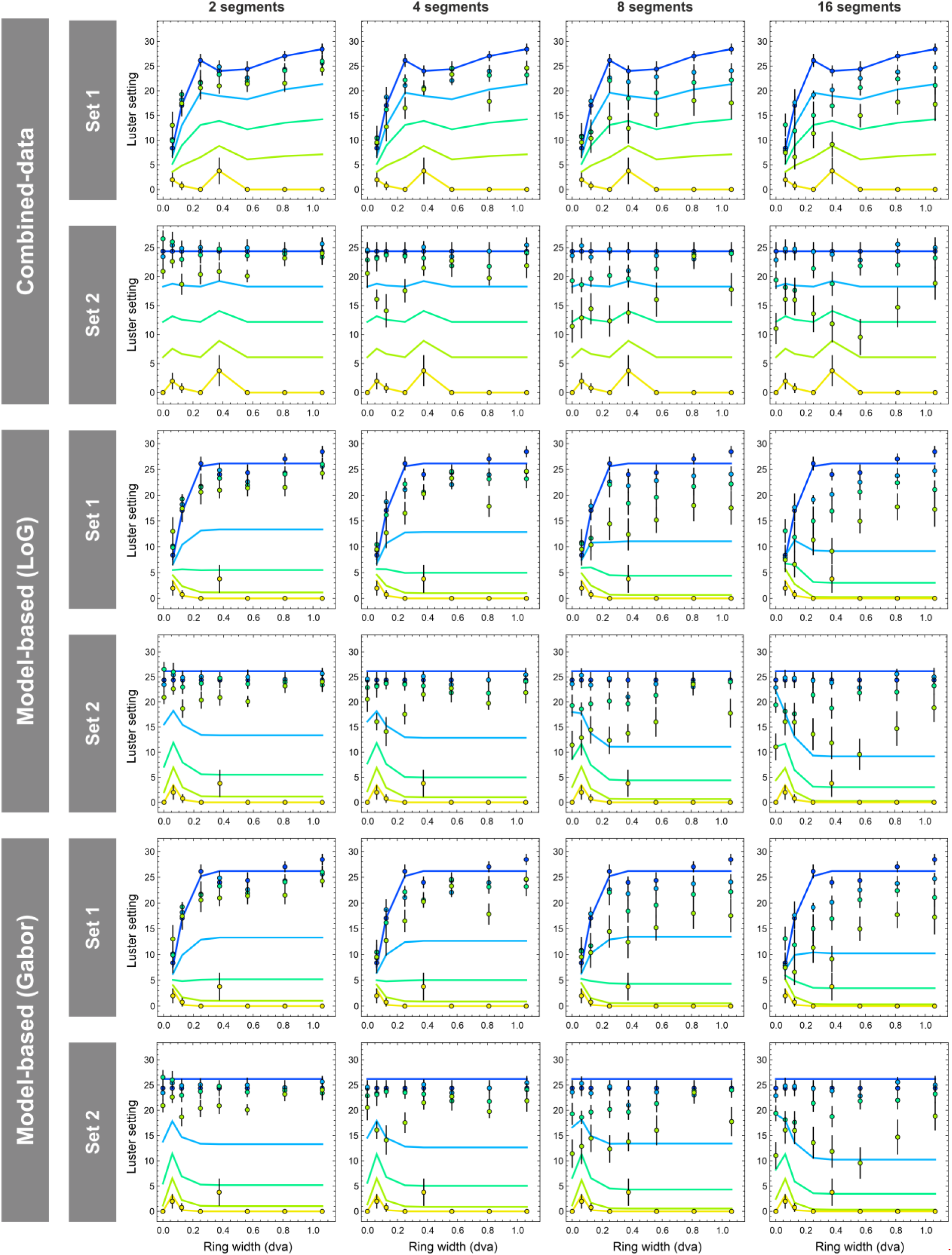
Empirical luster settings (colored disks) and corresponding predictions (colored lines) for group **A-**, separately for the three different methods used to calculate the predictions for the balanced integration process, that is, the combined-data method (top panel), and the two model-based methods, which are either based on an LoG kernel (middle panel) or a Gabor kernel (bottom panel). Each panel shows the results for the two different stimulus sets (rows), separately for the different numbers of ring segments (columns) and the different ring proportions (differently colored symbols and lines). The error bars represent SEM in both directions.

## Appendix E ANOVA results for the homogeneity rating data

**Table E1:** Top: Results of a three-way ANOVA performed on the homogeneity ratings for the combined datasets of groups **A+** and **A-** (top panel) based on the factors “group” (A+ and A-), “number of segments” (with the levels 1, 2, 4, 8, and 16), and “ring proportion” (with the levels 0.0, 0.25, 0.5, 0.75, and 1.0). Middle and bottom: Results of a two-way ANOVA performed on the homogeneity rating data for the mixed-ring conditions only, separately for group **A+** (middle) and **A-** (bottom). In these cases, the “number of segments” variable was tested only with the levels 2, 4, 8, and 16, and the “ring proportion” variable only with the levels 0.25, 0.5, and 0.75.

| **A+ & A-** |  |  |  |  |  |
| --- | --- | --- | --- | --- | --- |
| **Source** | **df** | **SS** | **MS** | **F** | **Sig.** |
| Group | 1 | 1788.55 | 1788.55 | 518.611 | p < 0.0001 |
| Segments | 4 | 2640.49 | 660.122 | 191.41 | p < 0.0001 |
| Proportion | 3 | 1264.84 | 421.612 | 122.251 | p < 0.0001 |
| Group * Segments | 4 | 106.323 | 26.5808 | 7.70741 | p < 0.0001 |
| Group * Proportion | 3 | 516.203 | 172.068 | 49.893 | p < 0.0001 |
| Proportion * Segments | 6 | 251.568 | 41.928 | 12.1575 | p < 0.0001 |
| Group * Proportion * Segments | 6 | 46.285 | 7.714 | 2.237 | p = 0.037 |
| Error | 4360 | 15036.5 | 3.44873 |  |  |
| Total | 4387 | 21650.7 |  |  |  |
| **A+** |  |  |  |  |  |
| **Source** | **df** | **SS** | **MS** | **F** | **Sig.** |
| Segments | 3 | 613.54 | 204.51 | 84.16 | p < 0.0001 |
| Proportion | 2 | 152.79 | 76.396 | 31.438 | p < 0.0001 |
| Segments * Proportion | 6 | 83.118 | 13.853 | 5.701 | p < 0.0001 |
| Error | 2147 | 5217.29 | 2.43 |  |  |
| Total | 2158 | 6066.74 |  |  |  |
| **A-** |  |  |  |  |  |
| **Source** | **df** | **SS** | **MS** | **F** | **Sig.** |
| Segments | 3 | 245.835 | 81.945 | 18.014 | p < 0.0001 |
| Proportion | 2 | 1632.11 | 816.057 | 179.394 | p < 0.0001 |
| Segments * Proportion | 6 | 214.736 | 35.789 | 7.868 | p < 0.0001 |
| Error | 1974 | 8979.65 | 4.54896 |  |  |
| Total | 1985 | 11072.3 |  |  |  |

